# Metabolic Buffer Analysis Reveals the Simultaneous, Independent Control of ATP and Adenylate Energy Ratios

**DOI:** 10.1101/2020.02.19.957035

**Authors:** Edward J. Hancock, James Krycer, Jordan Ang

**Affiliations:** School of Mathematics and Statistics, The University of Sydney, NSW, 2006, Australia; School of Life and Environmental Sciences, The University of Sydney, NSW, 2006, Australia; Charles Perkins Centre, The University of Sydney, NSW, 2006, Australia; Department of Chemical and Physical Sciences, The University of Toronto, ON, L5L1C6, Canada; Department of Immunology, The University of Toronto, ON, L5L1C6, Canada

## Abstract

Determining the underlying principles behind biological regulation is important for understanding the principles of life, treating complex diseases, and creating *de novo* synthetic biology. Buffering - the use of reservoirs of molecules to maintain molecular concentrations - is a widespread and important mechanism for biological regulation. However, a lack of theory has limited our understanding of its roles and quantified effects. Here, we study buffering in energy metabolism using control theory and novel buffer analysis. We find that buffering can enable the simultaneous, independent control of multiple coupled outputs. In metabolism, adenylate kinase and AMP deaminase enable simultaneous control of ATP and adenylate energy ratios, while feedback on metabolic pathways is fundamentally limited to controlling one of these outputs. We also quantify the regulatory effects of the phosphagen system - the above buffers and creatine kinase - revealing which mechanisms regulate which outputs. The results are supported by human muscle and mouse adipocyte data. Together, these results illustrate the synergy of feedback and buffering in molecular biology to simultaneously control multiple outputs.

## Introduction

Determining the design principles behind biological regulation is important for understanding the principles of life, treating complex diseases, and creating *de novo* synthetic biology [20]. A significant amount of research into this topic has focussed on feedback regulation [2, 8, 9, 18, 20, 34, 35, 36, 43], a ubiquitous mechanism in biology that acts via biological actuators, such as regulated enzymes [17, 38]. But there is a crucial open challenge to determine the underlying principles for biological regulation beyond feedback [20]. Buffering—the use of reservoirs of molecules to maintain molecular concentrations—is another widespread mechanism for biological regulation [17, 38]. Despite its importance, buffering has received considerably less attention than feedback. In recent work, we studied the interaction of feedback and buffering, where we found that synergies between them are often critical for biochemical regulation [17, 16]: Feedback regulates ‘slow’ disturbances while buffering regulates ‘fast’ disturbances and stabilises feedback.

However, we do not know all of the underlying principles of buffers and have a limited methodology for quantifying their effects. Our recent findings describe the principles where there is buffering of a single regulated output. While important, they do not fully describe cases where buffers affect multiple outputs, such as in energy metabolism [5]. Also, the familiar treatment for quantifying the effects of buffering is limited to closed systems (with mass and energy isolated from surroundings), whereas *in-vivo* biological systems are fundamentally open systems [20]. Further, the methodology is heavily focussed on pH buffering, while biological systems are predominantly non-pH buffers. Thus, there is a requirement for novel theory relating buffering to multiple outputs, biochemical kinetics, and the quantification of regulation in open systems.

### Box 1: Multivariable Control in an Energy Metabolism Context

In control theory, the simultaneous control of multiple coupled outputs is referred to as ‘multivariable control’. It is an important topic that has been extensively studied in technology and other non-biological contexts [39], but not in biological ones. However, the control of multiple coupled outputs is also important in biology, such as in energy metabolism. In this paper, we provide a foundational case study for multivariable control in cellular biology.

To help describe multivariable control in an energy metabolism context, we can compare the control of an aircraft’s (rotational) direction with that of the adenine nucleotide energy environment of a cell. We need to use three variables to describe the direction of an aircraft; roll, pitch and yaw (see below figure [41]). Without all three, we don’t have a complete picture of the aircraft’s direction To describe the adenylate energy environment of a cell, we similarly need to use three variable as there are three molecules involved (ATP, ADP and AMP). The important outputs are ATP, the ATP/ADP ratio, the ATP/AMP ratio and the energy charge (see introduction for descriptions), noting that one is redundant when all are used in combination.

To control the direction of an aircraft or energy environment of a cell, the three outputs all need to be simultaneously controlled. For an aircraft, the rudder (yaw), ailerons (roll), and elevators (pitch) are the regulatory mechanism that carry out the control of each of these outputs. For energy metabolism, the energy environment is controlled by the reaction rates of the metabolic pathways as well as creatine kinase, adenylate kinase and AMP deaminase.

In the below analysis, we first show that these buffers are required to simultaneously and independently control the three coupled outputs. We then quantitatively characterise the buffers to show which regulatory mechanisms regulate which outputs.

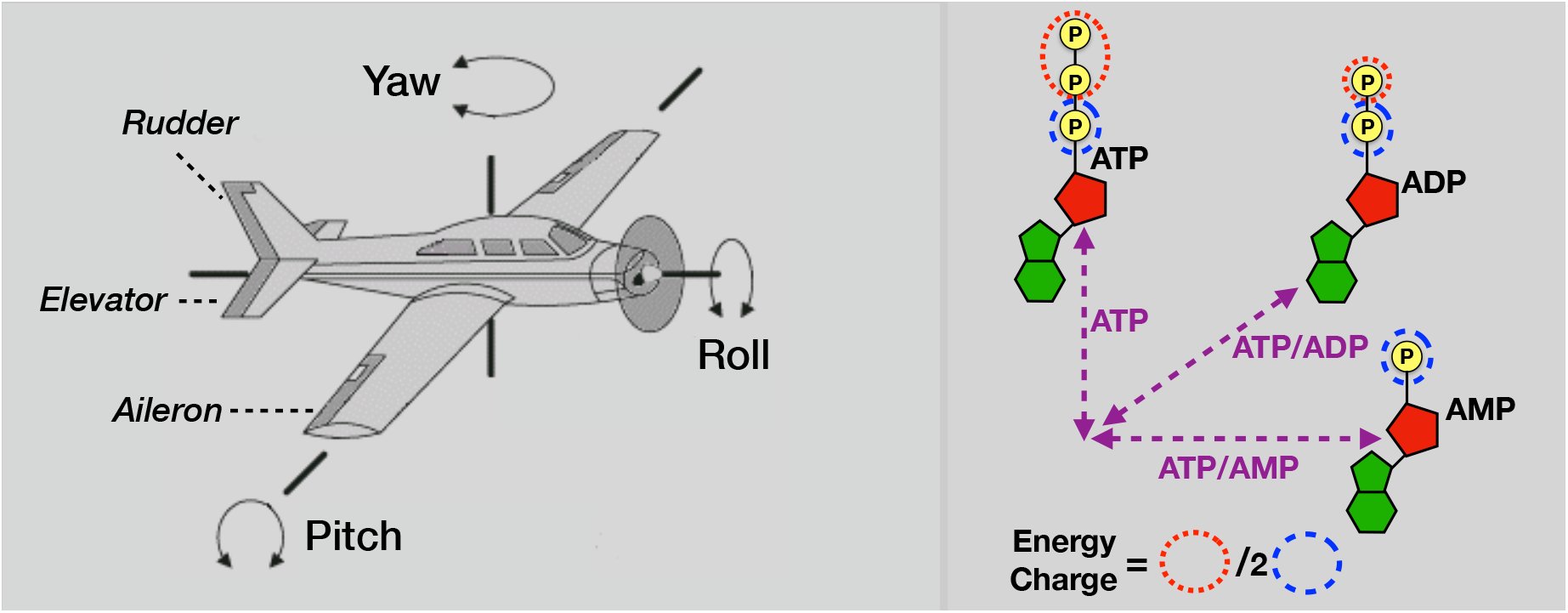

To understand regulatory roles in the case of energy metabolism, we need to know what outputs are regulated by which mechanisms from what types of disturbances [4, 17]. Disturbances, such as changing energy demand, cause deviations in the outputs due to a mismatch of ATP demand and supply, while the regulatory mechanisms reduce these deviations. Studies focus on the following key outputs: ATP concentration, which is the cellular energy currency; and the ATP/ADP and ATP /AMP concentration ratios, which are related to Gibbs free energy, i.e., the ability of a cell to do work (also see Box 1). These quantities are analogous to charge (ATP) and voltage (free energy) on batteries. Studies also focus on the adenylate energy charge *(ATP* + 0.5*ADP*)/(*ATP* + *ADP* + *AMP*), an index used to measure the overall energy status of biological cells [5] (see figure in Box 1). A widespread view is that cells primarily regulate for the energy charge or related AMP-based outputs, which is based on the energy charge’s steady dynamics and the observation that signalling and allosteric feedback primarily regulates for these outputs [5].

Despite extensive studies on metabolic buffers, a lack of theory has limited our understanding of their roles and quantified effects. This limitation is particularly important for understanding the phosphagen system [6], a critical buffering system in energy metabolism that is composed of adenylate kinase (AK), AMP deaminase (AMPD), and creatine kinase (CK). In the phosphagen system, creatine kinase regulates for fast-changing disturbances and stabilises oscillations caused by glycolytic feedback [17, 42], while AK and AMPD are buffering enzymes that have a number of regulatory roles relating to different energy outputs [5, 25, 30]. AK catalyzes the regeneration of ATP from ADP and enables AMP-based feedback by also producing AMP [5, 25]. AMPD removes AMP and limits its accumulation [5]. It has been proposed that AMPD’s role is to regulate adenylate ratios [5]. Other studies propose that AK and AMPD buffering is intended to limit the accumulation of ADP in muscles [15, 14]. However, it is difficult to collectively interpret the phosphagen system’s many roles without understanding the underlying principles, or to gauge their relative importance without quantification.

Here, we study buffering in energy metabolism using control theory and novel buffer analysis for open systems. We find that buffering can enable the simultaneous, independent control of multiple coupled outputs, which is in addition to its known role of regulating ‘fast’ disturbances. In energy metabolism, we find that adenylate kinase and AMP deaminase enable the simultaneous control of ATP and the key adenylate energy ratios, whereas feedback on metabolic pathways is fundamentally limited to controlling one of these outputs. This result significantly differs from, but can also explain, the widespread view that cells primarily regulate for a single adenylate energy output. We also quantify the regulatory effects of the phosphagen buffering system to reveal which mechanisms regulate which outputs. We find that AK regulates the ATP/ADP ratio while AMPD regulates the ATP/AMP ratio and the energy charge. We find that creatine kinase is significantly more effective when the ATP/ADP ratio is highly sensitive to changes. The phosphagen system quantification is shown to match with human muscle and mouse adipocyte data, including with AK knockout data. Together, these results illustrate the synergy of feedback and buffering in cellular biology to simultaneously control multiple outputs.

We use two separate but complementary methods in the paper: in section 2 we qualitatively study the control of multiple outputs using control theory, and in section 3 & 4 we quantify the regulatory effect of the phosphagen system using novel buffer analysis.

## 1 Background: Buffer Analysis for a Single Output

Buffering, the use of reservoirs of molecules to maintain molecular concentration, is a widespread mechanism for biomolecular regulation. In recent work, we found that the synergy between buffering and feedback is often critical for biochemical regulation [17, 16]. In this section, we provide a background on buffering for systems with a single output. We use a minimal model to help explain the quantification metrics we use for buffering, noting that the metrics described in this secti on apply generally.

Consider the model of a single regulated species that is buffered (see Figure 1)

**Figure 1:**
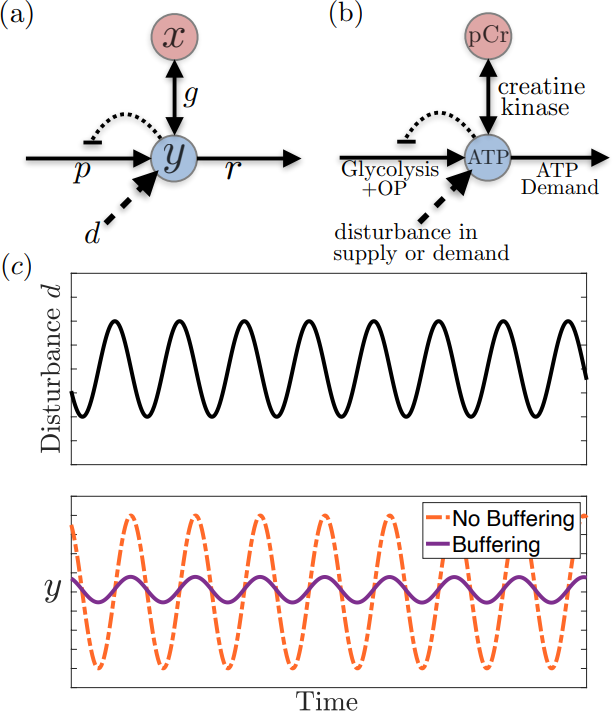
(a) Buffering and feedback schematic for a single output. *x* and *y* are the concentrations of the buffer and regulated species, respectively. *p* and *d* are the the production and disturbance rates of the regulated species. (b) An example with creatine kinase (CK) buffering; pCr is creatine phosphate, Cr is creatine and OP is oxidative phosphorylation. (c) Simulations of (1) showing buffering regulating a ‘fast’ oscillating disturbance.

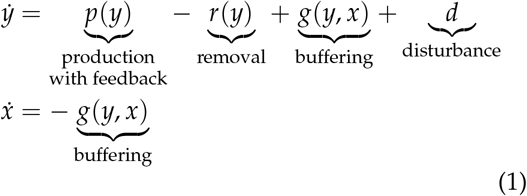

where *y* is an output concentration being regulated, *x* is the concentration of the buffering species, *p* is the production rate of y, *r* is the removal rate of *y, g* is the net flux through the buffering reactions, and *d* is a disturbance to the system. Incorporation of feedback is represented by the *y* dependence of production. The nominal steady state (when *d* =**0**) occurs when production matches degradation *(p(y) = r*(*y*)) and the buffer is at equilibrium *(g(y,x) =* 0). An illustrative example with creatine kinase buffering can be found in Figure 1.

To quantify the regulatory effects of systems with feedback and buffering, we use two key measures of regulation: the sensitivity function and the buffer equilibrium ratio [17]. The sensitivity function quantifies the overall regulatory effectiveness produced by all regulatory mechanisms present (e.g., both feedback and buffering), while the buffer equilibrium ratio is a metric specific to buffering. The latter is independent from other forms of regulation and from the disturbance itself (see SI 1.4 for analysis of a simple example).

The buffer equilibrium ratio compares the change in an output concentration, *y*, to the change in the concentration of a buffering species, *x*, when the buffering reactions are at (quasi) equilibrium [17] (see Figure 1). This is written as

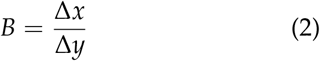

where 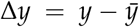 and 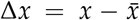 are the deviations of *y* and *x* from their undisturbed steady states (“set points”) 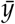 and 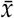 i.e. when there are no disturbances (see SI1.1). Thus, *B* represents the degree to which the effects of the disturbance have been absorbed by the buffer vs permeated through to the system output. The metric, which describes a relationship at (quasi) equilibrium, is important for analysis of both temporal dynamics and steady state of a system (see below). For small disturbances, *B* can be locally approximated about the system set point (see SI1.2). An equivalent approximation is used to define the well-known “buffer capacity” metric for pH regulation in closed systems (see SI3), and for steady-state sensitivity analysis as commonly applied in systems biology [34].

If we linearise the simple case in (1) and assume that the buffering reactions rapidly reach (quasi) equilibrium then the model can be simplified to (see SI1.1)

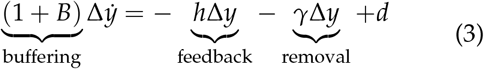

Where 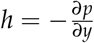 is the linearised feedback gain and the buffer concentration is determined by Δ*x = B*Δ*y*.

We can observe in (3) that buffering reduces the rate of change of the output Δ*y* by a factor of (1 + *B*), which attenuates the effect of ‘fast’ disturbances (see Figure 1)[17]. In contrast, buffering in (3) does not affect steady state disturbances, where any changes in *B* have no effect on (3) when 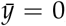. Thus one use of *B* is to measure a buffer’s ability to regulate ‘fast’ disturbances [16,17], where large *B* implies improved regulation. Buffering can also stabilise oscillations caused by feedback, such as those from delay in the feedback mechanism [17]. B also has an important control-theory interpretation in terms of derivative feedback; the net flux through the buffering reactions is 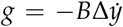 (see SI1.3) [16,17]. In technology, derivative feedback is a critical component of PID (proportional + integral + derivative) controllers, a ubiquitous form of feedback controller [4].

It is important to distinguish *B* from the familiar thermodynamic equilibrium constant, *K:* the latter relates (zero-disturbance) steady-state concentrations directly (e.g., 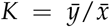), while *B* relates their (non-zero disturbance) deviations from steady state.

The sensitivity function is a normalised measure of the change in a system output, y, due to a disturbance, *d*. Again, it is affected by all internal regulatory mechanisms present. The sensitivity function is written as

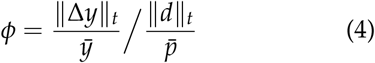

where ‖Δ*y*‖_*t*_ represents the size of deviation Δ*y* accumulated over time, ‖*d*‖_*t*_ represents the size of disturbance *d* accumulated over time, 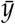 is the set point, and 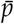 is the production rate of *y* at the set point. The exact form of ‖*d*‖_*t*_ and ‖Δ*y*‖_*t*_ depends upon the type of disturbance. For the example in Figure 1 with an oscillating disturbance, both would measure the amplitude of the oscillations.

The sensitivity function (4) is closely related to commonly used sensitivity analysis in systems biology for studying steady states [34], noting that the function used here can incorporate temporal dynamics as well as steady-state analysis.

The reader may find it instructive to compare and contrast the analysis of pH buffering with open-system metrics introduced in Section SI1 to familiar metrics commonly used to analyze closed-system, which can be found in SI3 (noting that this comparison is original to this paper).

## 2 Buffering can Enable the Control of Multiple Outputs

In this section, we ask what concentrations and outputs can feedback and buffering control? This question is well suited to controllability analysis from control theory [4]. Controllability determines the ability for regulatory mechanism inputs (e.g. enzyme activity) to change the system states (concentrations). Determining the actions of regulatory mechanisms contrasts to determining the outputs the mechanism responds to or senses. For example, a pathway enzyme may respond to a change in AMP by acting on ATP production, where controllability characterises the effect on ATP and not the sensing of AMP.

We first use a minimal model to illustrate general principles of controlling multiple outputs with buffering and feedback. We then apply the general principles to demonstrate the phosphagen system’s role in controlling multiple adenylate energy outputs.

### 2.1 Control of Multiple Outputs in a Minimal Model

To study the role of buffering, we use a modification of the minimal model in (1) to include two outputs with buffering on the second output. However, in this section we do not assume that the buffer rapidly reaches equilibrium. Consider (see Figure 2a)

**Figure 2:**
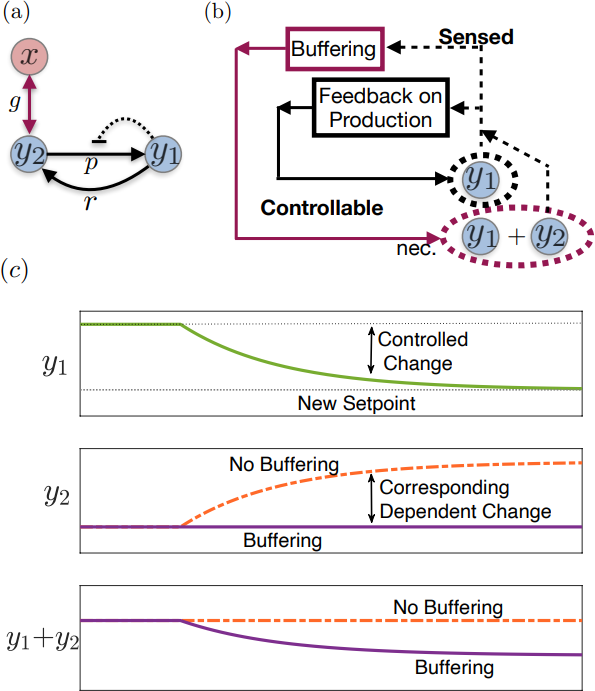
(a) Buffering and feedback schematic for multiple outputs - modified from Figure 1a to include two outputs and a buffer on the second output. *y*_*1*_ and *y*_*2*_ are the regulated output species and *x* is the buffering species. (b) Representation of controllability of the system in (a) in terms of *y*_1_ and *y*_1_ *+ y*_2_. Buffering is shown to enable controllability of *y*_1_ + *y*_2_, for which it is necessary (nec.), but it may also help control *y*_1_ (not shown). (c) Simulations of (5) show buffering enabling the simultaneous, independent control of multiple outputs. The ideal case is shown where buffering perfectly compensates *y*_2_ for a change in *y*_1_.

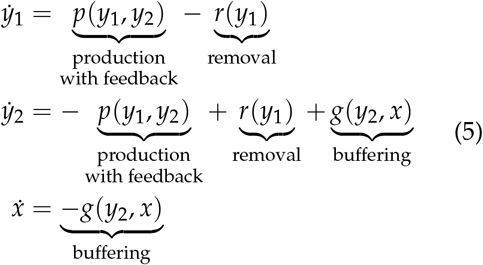

where *y*_1_ and *y*_*2*_ are concentrations of two output species being regulated, *x* is the concentration of the buffering species, *p* is the production rate of *y*_1_, *r* is the removal rate of *y*_1_, and *g* is the net flux through the buffering reactions. Now

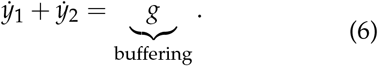

It is possible for feedback on production to control the concentration of *y*_1_. However, without buffering *(g =* **0**) we have 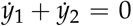 which results in constant *y*_1_ + *y*_*2*_ *= D* [37,31]. This implies that if the concentration of *y*_1_ is controlled to any particular concentration, then *y*_*2*_ = *D* − *y*_1_ is also determined and thus *y*_2_ is dependent upon *y*_1_ or vice versa (see Figure 2c). Therefore, without buffering then feedback cannot independently control the concentrations of *y*_1_ and *y*_2_. The total *y*_1_ + *y*_2_ is required to be controlled in order to simultaneously and independently control both *y*_1_ and *y*_2_, or combinations of the two e.g. *y*_1_ and *y*_1_/ *y*_2_*-*Buffering can control *y*_1_ + *y*_2_ and thus enable the concentrations of *y*_1_ and *y*_2_ to be independently controlled via the net buffering flux *g* (see Figure 2c). A representation of the controllability can be seen in Figure 2b.

We can note that the buffer in model (5) (see Figure 2) can take the role described in this section, but it can also attenuate fast disturbances that act on *y*_*2*_, as described in Section 1. Also, although not the focus of this paper, the above regulatory principles are relevant for biological systems other than metabolism, such as signalling and gene regulatory networks.

### 2.2 The Phosphagen System

We next apply these principles to illustrate the role of the phosphagen system [6] in controlling ATP, ADP and AMP This system is represented by the buffering reactions

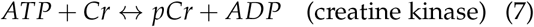

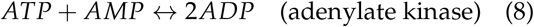

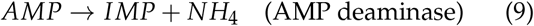

where ATP, ADP and AMP are the adenine nucleotides used for energy, pCr is creatine phosphate, Cr is creatine, and IMP is inosine monophosphate. IMP can be regenerated to form AMP in separate reactions. The right hand side of (7) and the left hand side of (9) can also include *H*^+^, but we do not include them here both for simplicity and so that we can later use standard definitions of equilibrium constant *K*.

To study controllability in the phosphagen system, we first look at the effect of adenylate kinase without AMP deaminase. We then use analysis of this subsystem to study controllability of the whole system with AMPD - a form of mathematical induction.

To determine the effect of the adenylate kinase (without AMPD) on controllability, we study the ATP and ADP concentrations and incorporate metabolic pathways that are controlled by various form of feedback (see Figure 3a and SI2 for analysis). We ignore any reactions that cannot be regenerated without AK (e.g. ATP→AMP). This case is very similar to the above minimal model if we set *y*_1_ = *ATP* and *y*_*2*_ = *ADP*. As above, feedback on metabolic pathways can control the concentration of ATP, but ATP+ADP is constant without the buffer AK and so feedback alone cannot independently control both ATP and ADP. Similarly, creatine kinase also has no ability to change the total ATP+ADP (see SI2). Adenylate kinase ensures system controllability by varying the total ATP+ADP (see Figure 3c for graphical representation). This controllability enables the independent control of ATP and ADP, or combinations such as ATP and ATP/ADP (also see SI2 for equivalent analysis of polyphosphate kinase).

**Figure 3:**
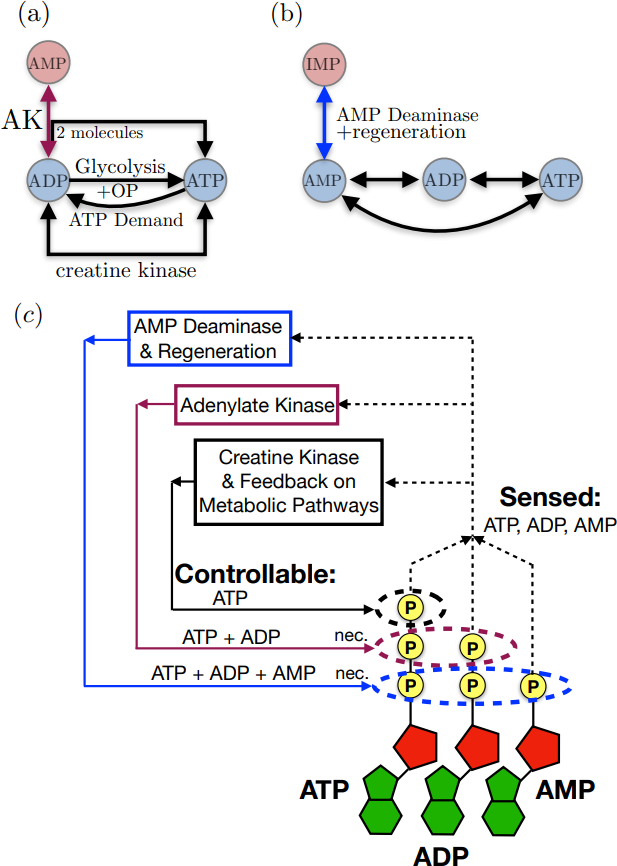
(a) Buffering schematic for Adenylate Kinase (AK) and Creatine Kinase - modified from Figure 2a (see (16) for reactions). (b) Buffering schematic for AMP Deaminase and regeneration (see (17) for reactions). (c) Representation of the controllability for the phosphagen system and feedback on metabolic pathways. AK and AMPD may control other outputs than those illustrated, where only those necessary (nec.) for controllability are shown.

While AK enables independent control of two outputs, to study the control of ATP, ADP and AMP we also need to incorporate AMPD and regeneration (see Figure 3b and SI2 for analysis). We ignore any processes that cannot be regenerated via resynthesis of IMP. If necessary, other regeneration reactions can be incorporated in the models as disturbances, rather than regulatory mechanisms, without changing the results e.g. production of AMP from the degradation of RNA. In this model, feedback and AK can control ATP and ADP, but without AMP deaminase ATP+ADP+AMP is constant and thus unable to be controlled. Similar to above, AMP deaminase can vary the total ATP+ADP+AMP and thus can enable the independent control of ATP, ADP and AMP or various combinations.

Thus we can observe that adenylate kinase and AMP deaminase enable the simultaneous, independent control of ATP, ADP and AMP, which is equivalent to enabling the simultaneous, independent control of ATP, ATP/ADP and ATP/AMP (see SI2). In contrast, the system with feedback on metabolic pathways but without buffering is not controllable and is fundamentally limited to only controlling one of these outputs.

We can also take an alternative approach to the one used here and treat AK buffering as part of the metabolic system rather than as a control mechanism. For this case, if we ignore AMPD then we have the conserved total *ATP* + *ADP* + *AMP = M* along with the AK equilibrium constant 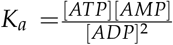, which result in three outputs and two dependencies. Thus we can observe that feedback is limited to controlling one independent output. However, without treating the buffer as a control mechanism then we do not observe the role of the adenylate kinase in independently controlling multiple outputs.

## 3 Quantifying Buffering

While the controllability analysis in the last section *shows* what the regulatory mechanisms can and cannot act on, it provides a qualitative result and does not show which outputs particular mechanisms are regulating, or by how much. For this question, we need to quantify the regulatory effects of energy buffers. In this section, we quantify the effects of creatine kinase and adenylate kinase buffering by determining the buffer equilibrium ratio of each. The reader may recall that this ratio is specific to the individual buffer and can measure each buffer’s ability to regulate ‘fast’ disturbances. We will see in Section 4 that the buffer equilibrium ratio of adenylate kinase is also important for quantifying the regulation of multiple energy outputs.

### 3.1 Creatine Kinase

We first determine the buffer equilibrium ratio for creatine kinase buffering. We find that creatine kinase is most effective at high *K* and high ATP/ADP ratios (or low K and low ATP/ADP ratios). We also show that the effectiveness of creatine kinase is reduced in human muscles by adenylate kinase. However, this is offset by the boost to the effectiveness of creatine kinase from varying equilibrium constant K (via changing pH). We show that the theory matches experimental data for human muscle during and after exercise.

The buffer equilibrium ratio for the creatine kinase reaction described in equation (7) is

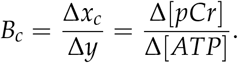

If we initially assume that the equilibrium constant *K*_*c*_ and other parameters are fixed then (see SI4.2)

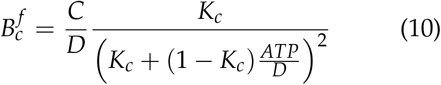

where *x*_*c*_ = *[pCr], y = [ATP], C =* [*pCr]* + [*Cr*], *D* = *[ATP]* + *[ADP]*, the equilibrium constant is *K*_*c*_ *= [ATP][Cr]/(\pCr][ADP])*, and C, D and *K*_*c*_ are the fixed parameters. We can observe that the buffer equilibrium ratio 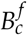 in (10) increases linearly with *C*/*D*, but the relationship between 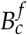 *K*_*c*_ and *ATP/ D* is more complicated.

In Figure 4, we can see that there are two regions with large 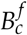 and consequently where pCr is most effective at regulating ‘fast’ disturbances. These regions occur where IQ is small together with small ATP/ADP, and where *K*_*c*_ is large together with large ATP/ADP, noting that ATP /ADP is large when ATP/D is close to 1. Outside of these regions 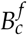 is significantly smaller and consequently the buffer is less effective.

**Figure 4:**
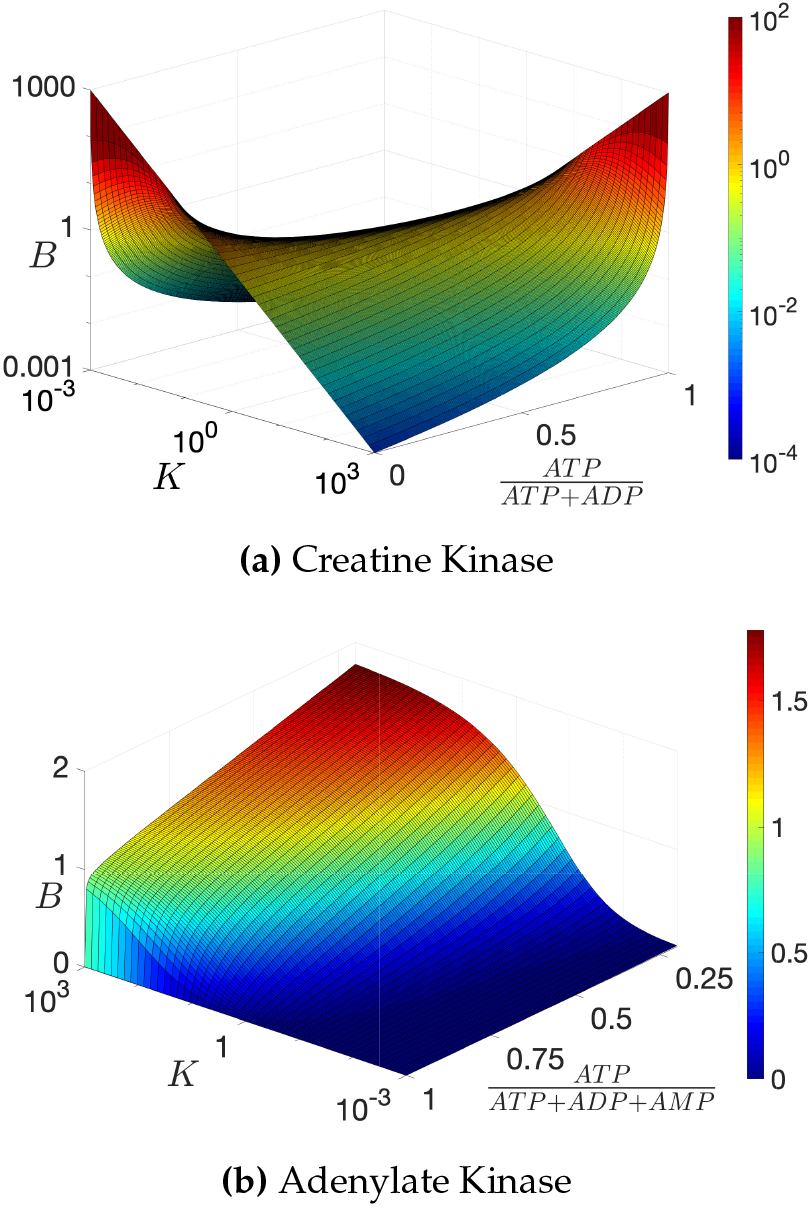
Buffer Equilibrium Ratios for CK and AK, which quantify their buffering effects under different conditions. CK has normalisation C/D = 1 where C = [*pCr*] + [*Cr*] and D = [*ATP*] + [*ADP*].

In calculating *B*_*c*_, we also need to take the varying equilibrium constant *K*_*c*_. and D = *ATP* + *ADP* into account. For example, during exercise the use of ATP decreases intracellular pH, which increases the value of *K*_*c*_. Similarly, parameter *D* decreases as ATP levels decrease due to adenylate kinase converting ADP to AMP. For varying *K*_*c*_ and D, the buffer equilibrium ratio of pCr is (see SI4.2)

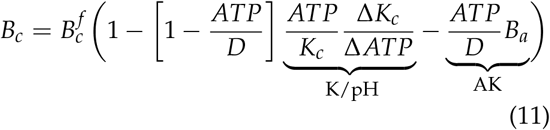

Where 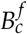 is given in (10), *B*_*a*_ = Δ*D/*Δ*ATP* is the rate of change of D = *[ATP]* + [ADP] with respect to *ATP* (also see section 3.2), Δ *K*_*c*_ */ΔATP* is the rate of change of equilibrium constant *K*_*c*_ with respect to ATP, and total creatine C = *Cr* + *pCr* in (10) is assumed constant. It can be observed that the second term is due to changing *K*_*c*_ (via changing pH) while the third term is due to adenylate kinase.

Under high energy demand situations like exercise, *K*_*c*_ typically increases when ATP drops (via decreasing pH), which shifts reaction (7) to the left back towards ATP and increases the counteracting effect of buffering (↑ *B*_*c*_). In contrast, AK removes more ADP when ATP decreases, which shifts reaction (7) to the right a way from ATP and reduces the counteracting effect of buffering (↓ *B*_*c*_).

We can compare the above theory to experimental data from literature, where Figure 5 shows values of buffer equilibrium ratio *B*_*c*_ for human muscles during and after exercise (see SI4.2.3 for calculation methods). It can be seen that the theoretical values of *B*_*c*_ without AK or varying *K*_*c*_ are not consistent with experimental values, but match with experimental values when these are included (also see SI4.2).

**Figure 5:**
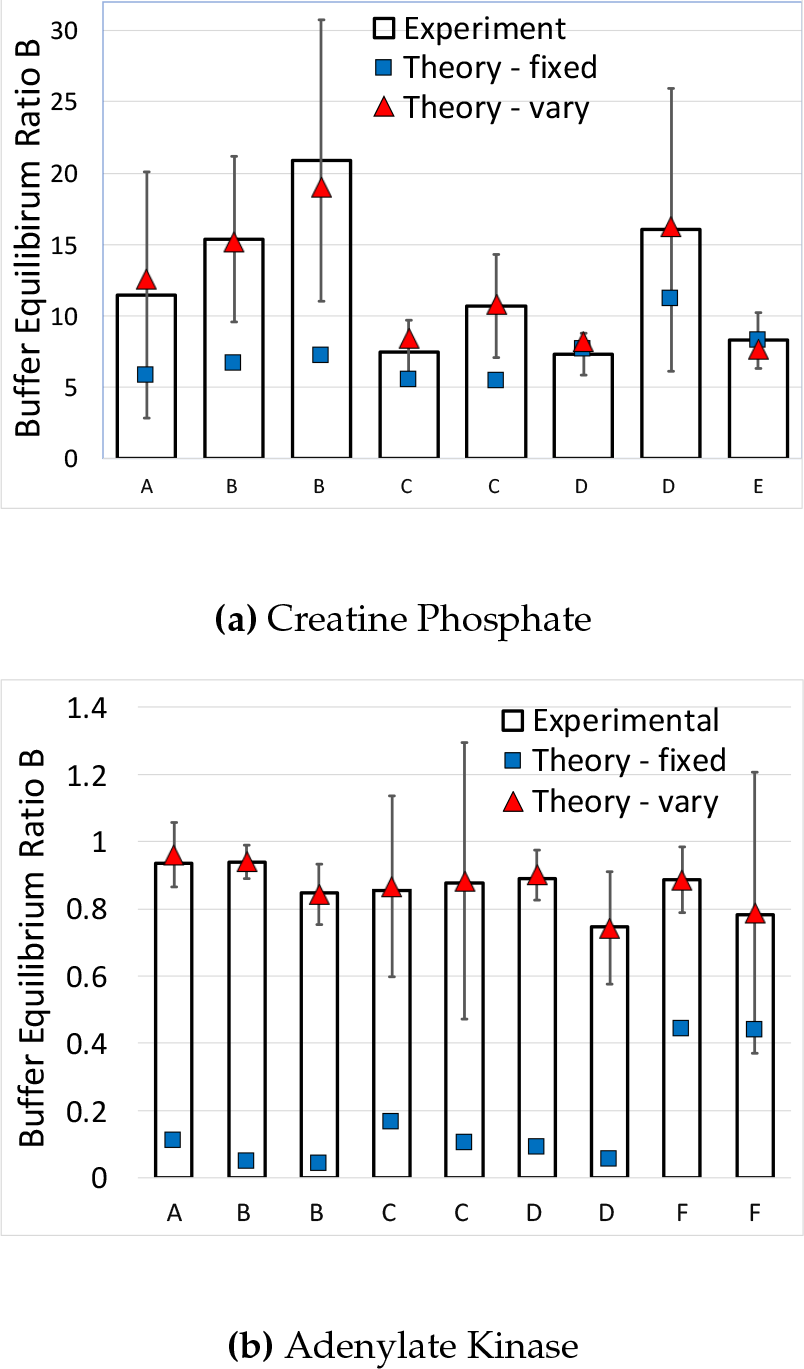
Comparison of Experimental and Theoretical Buffer Equilibrium Ratios with fixed and varying parameters. A-E show human muscle data and F shows mouse adipocyte data (mean ± SE) - See Appendix.

We can observe that the experimental value of *B*_*c*_ for CK ranges from 7.3 to 21. The metric 1/(1 + *B*_*c*_) can act as a proxy for the attenuation of fast disturbances by CK (see SI1.4). Thus the results infer that CK reduces deviations from fast disturbances by an order of magnitude.

### 3.2 Adenylate Kinase

In this section, we quantify the buffer equilibrium ratio of adenylate kinase under different conditions. We find that AK typically has a buffer equilibrium ratio that is slightly less than one, which implies that its importance for regulating for ‘fast’ disturbances is secondary and significantly smaller than creatine kinase. We also show that AMPD increases the effect of AK, although the increased ratio is still slightly less than one in the experimental data. We show that the theory matches experimental data for human muscle and mouse adipocytes.

To study the buffering properties of the adenylate kinase reaction described in Equation (8), we set the output as *y = [AT P]* and the buffer or reservoir as *x*_*a*_ = *D* = *[ATP]* + [*ADP*]. If we had instead set *ADP* as the buffer then the ATP hydrolysis *ATP → ADP* would be a buffering reaction alongside reaction (8). By setting *x*_*a*_ *=* D, then ATP hydrolysis does not simultaneously remove *y* and produce *x*_*a*_ and so is independent of the buffering reactions.

The buffer equilibrium ratio for adenylate kinase is

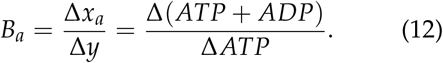

If we initially assume that the equilibrium constant *K* and other parameters are fixed then (see SI4.3)

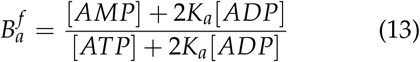

Where 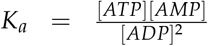 and the adenine nucleotide total A = *ATP* + *ADP* + *AMP* is a fixed parameter. Equation (13) can also be written in a more complicated form as an explicit function of two independent inputs *Ka* and *ATP/A* (see SI4.3). A related result for the alternate purpose of determining changes in AMP to changes in ATP can be found in [34, 11] - changes in AMP correspond to changes in ATP + ADP if A is fixed (see above) but not if A is varying (see below).

We can observe in Figure 4 that buffer equilibrium ratio 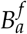 is higher for higher *K*_*a*_ values. However, 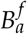 is not large (compared to creatine kinase) at high *ATP/ A* and *K*_*a*_, with a maximum at or near one. Although not shown in the figure, 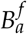 can become large at very low energy charges where *AMP* > *ATP* (see SI4.3).

We also need to take AMPD and corresponding regeneration into account by incorporating varying *A*. For this parameter-varying case, the buffer equilibrium ratio of AK is (see SI4.3)

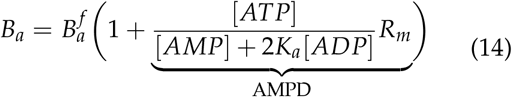

where *R*_*m*_ = Δ *A*/ Δ *ATP* is the rate of change of A with respect to *ATP* due the removal of AMP by AMP deaminase and the corresponding AMP regeneration (see SI4.3 for the effect of changing *K*_*a*_).

AMPD removes AMP as ATP decreases, which shifts reaction (8) to the left back towards ATP and increases the counteracting effect of buffering (↑ *B*_*c*_).

We can compare the theoretical buffer equilibrium ratios to human muscle and mouse adipocyte experimental data. Figure 5 shows values of theoretical and experimental *B*_*a*_. It can be seen that the theoretical values of *B*_*a*_ from (3) are not consistent with experimental values when AMPD is not incorporated, but match with experimental values when it is (also see SI4.3). It can be noted that the adipocyte data also requires the assumption that *K* varies. The effect of AMPD on AK can be seen in Figure 5 to be much larger in muscle than in adipocytes, but that the buffer equilibrium ratio is just below one in both cases.

Figure 5 shows that the AK buffer equilibrium ratio *B*_*a*_ is slightly less than one in human muscle during/after exercise, while the pCr ratio is *B*_*c*_ *=* 7.3 − 21 during/after exercise. As the total buffer equilibrium ratio for ATP is *B*_*TOT*_ = *B*_*c*_ +B_*a*_ [17], we can observe that AK is much less important than pCr in regulating for ‘fast’ disturbances.

## 4 Quantifying Ratio Regulation

In this section, we quantify the overall effect of the phosphagen system to regulate key energy outputs: and the energy charge. We find that AK regulates the ATP/ADP ratio while AMPD regulates the ATP/AMP ratio and the energy charge. We also find that AK worsens regulation of the ATP/AMP ratio and the energy charge, but this tradeoff is compensated for by AMPD. The reader may recall that the ATP/ADP and ATP/AMP concentration ratios are related to Gibbs free energy, i.e. the ability of the cell to do work, while the adenylate energy charge (*ATP* +0.5 *ADP)/(ATP* + *ADP* + *AMP)* is an index used to measure the overall energy status of biological cells [5]. We also show that the theory matches experimental data for human muscle and mouse adipocytes.

To quantify the regulation of the energy ratios, we determine the sensitivity of the energy ratios to changes in ATP. Determining these sensitivity functions enables a simple means of translating sensitivity function (4) for ATP into sensitivity functions for the energy ratios (see SI5.5), as well as being important is its own right e.g. it is analogous to relating the charge (ATP) and voltage (ratios) of a battery. We use ATP as a reference as the regulation metrics are most naturally written in terms of ATP concentrations e.g. disturbances are most naturally represented in terms of ATP demand or supply. We assume that AK is at (quasi) equilibrium and that AMPD and its corresponding regeneration are the primary means of interconversion between AMP and IMP.

The sensitivity of the ATP/ADP ratio, ATP/AMP ratio and energy charge to changes in ATP are (see SI5.1)

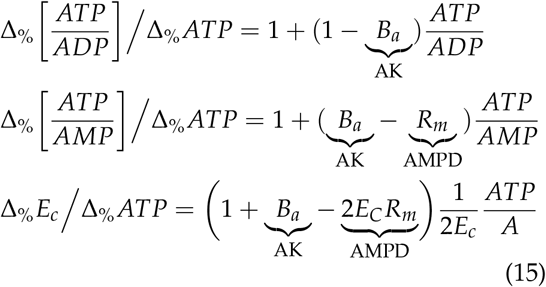

Where Δ_%_ *ATP* = Δ *ATP*/*ATP* is the percentage change in [*ATP*], 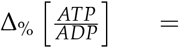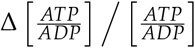, Δ_%_ *E*_*c*_ = Δ*E*_*c*_/*E*_*c*_ and 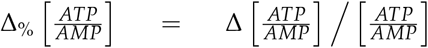 are the percentage change in the respective rations, *B*_*a*_ is defined in (12), *R*_*M*_ is defined below (14), A is defined below (13) and *E*_*c*_ = (*ATP*+0.5 *ADP*)/(*ATP* +*ADP*+*AMP*).

Equation (15) is plotted in Figure 6 and shows that at high ATP/ADP or ATP/AMP ratio then small changes in ATP can result in large changes in both the ATP/ADP and ATP/AMP ratios respectively (first reported in [27]). This effect occurs as small relative changes in ATP cause large relative changes in ADP and AMP The energy charge does not suffer from the same sensitivity effects as the other ratios (see Figure 4).

**Figure 6:**
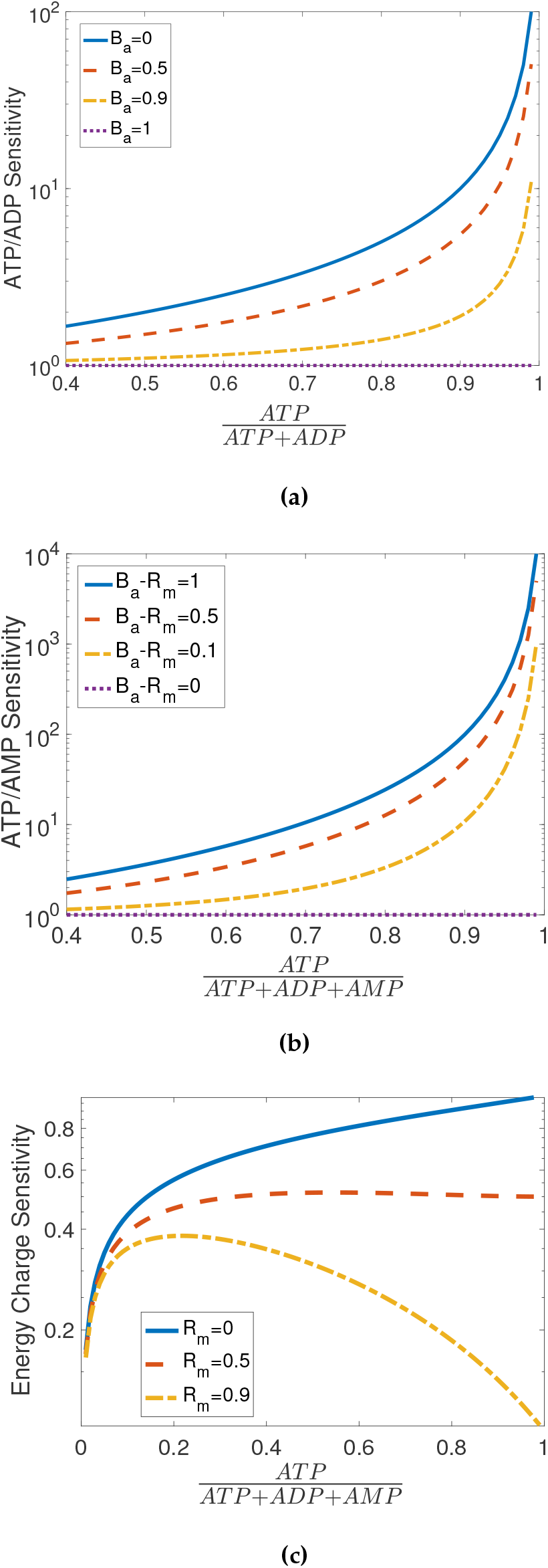
Sensitivity of Adenylate Energy Ratios to changes in ATP (as defined in (15)). These plots show the regulatory effect of AK (via *B*_*a*_) and AMPD (via *R*_*m*_) on different outputs.

Equation (15) and Figure 6 show that AK reduces the sensitivity of the ATP/ADP ratio to changes in ATP, reaching a minimum when *B*_*a*_ = 1. AK also creates a tradeoff as increasing *B*_*a*_ worsers the regulation of the ATP/AMP ratio. However, AMPD regulates the ATP/AMP ratio by lessening this sensitivity effect, reaching a minimum when *R*_*m*_= *B*_*a*_. The regulation of the energy charge is also worsened by AK and improved by AMPD. It can be noted that the sensitivity functions in (15) are not a function of creatine kinase, which through *K*_*c*_= [*ATP*][*Cr*]/([*PCr*][*ADP*]) sets the sensitivity between pCr/Cr and ATP/ADR rather than that between ATP and ATP/A DP.

This analysis effectively quantifies the regulation of the energy ratios into two forms. First, improving ATP regulation can be observed to improve regulation of the energy ratios, as reduced deviations in the former will reduce deviations in the latter (or vice versa). Through this general means, it is possible for improved regulation of ATP from feedback and pCr to also improve the regulation of the energy ratios (or vice versa). However, due to the sensitivity of the ratios under some conditions, ATP may be well regulated while the energy ratios are poorly regulated. In the second form of regulation, AK and AMPD can reduce the sensitivity of the individual energy ratios to changes in ATP, enabling simultaneous effective regulation of ATP and the ratios. These observations are consistent with Section 2, where AK’s and AMPD’s abilities to enable simultaneous, independent control of multiple outputs also enables simultaneous, tailored regulation of the different energy ratios. Interestingly, while pCr regulates ‘fast’ disturbances, AK and AMPD reduce the sensitivity of the energy ratios to any deviation in ATP. Thus AK and AMPD help regulate both ‘fast’ and ‘slow’ disturbances.

We can see in Figure 7 that the sensitivity functions match experimental data for human muscle and mouse adipocytes. The results in the figure show that on average AK reduces the ATP/ADP sensitivity by × 3.9/4.5 (muscle/adipose), AMPD reduces the ATP/AMP sensitivity by × 37.3/ × 12.8 (muscle/adipose) and AMPD reduces the energy charge sensitivity by × 6.1 / × 5.4 (muscle/adipose).

**Figure 7:**
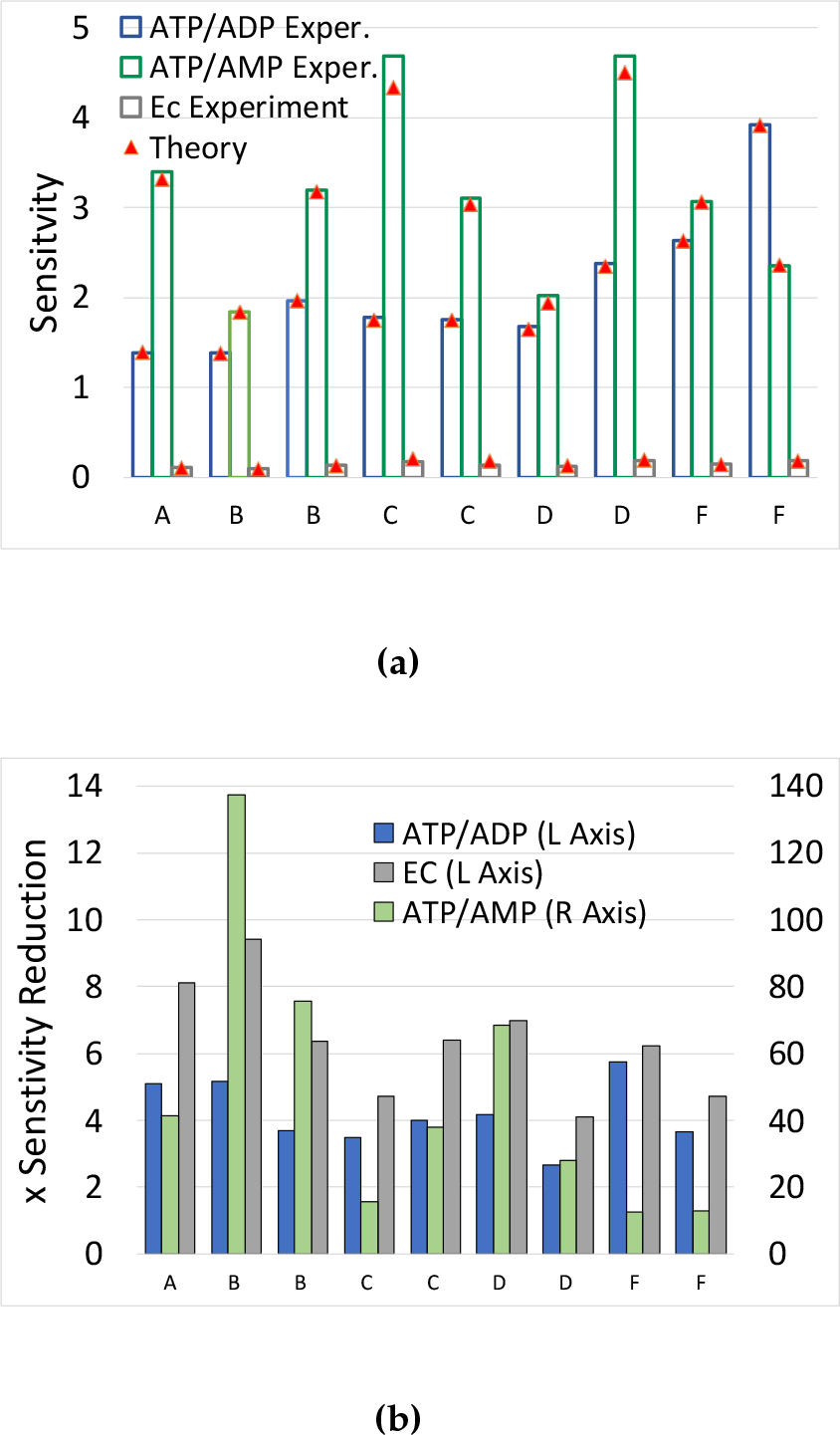
(a) Comparison of Experimental and Theoretical Sensitivity of ATP/ADP (left), ATP/AMP (centre) and the Energy Charge (right) to changes in ATP. (b) Sensitivity Reduction of ATP/ADP by AK (left), ATP/AMP by AMP deam. (centre) and the Energy Charge by AMP deam. (right). A-E show human muscle data and F shows mouse adipocyte data (See Appendix).

The inferred regulation of the ATP/ADP ratio by AK is supported experimentally by AK knock-out data, which resulted in a significant accumulation in ADP and worsening of the ATP/ADP ratio in skeletal muscle [14].

## 5 Discussion

The study of the regulatory roles of the phosphagen system illustrate an important general principle - buffering can enable the simultaneous, independent control of multiple coupled outputs. This role is in addition to its known role of regulating ‘fast’ disturbances and stabilising feedback. Different buffers can also carry out different roles. In the phosphagen system, AMPD and AK carry out the ‘independent control’ role, while CK and AK (to a much lesser extent) carry out the latter roles. In AK’s case, its ability to enable simultaneous control occurs due to its production and removal of AMP (and thus ATP+ADP), while its lesser second role occurs due to the regeneration of ATP. In an interesting difference between the roles, CK regulates ‘fast’ disturbances while AK and AMPD regulate both ‘fast’ and ‘slow’ disturbances.

How important is enabling simultaneous, independent control? In the case of energy metabolism, there are multiple outputs and their regulation that are widely regarded as biologically important (see intro). The coupling between these outputs implies that simultaneous and independent control is fundamental to this regulation of multiple outputs. Without simultaneous, independent control, the regulation of one output can interfere with the regulation of another e.g. AK without AMPD worsens the regulation of ATP/AMP and the energy charge. Also, due to the sensitivity of ratio outputs, it can be difficult to regulate a ratio on its own without independent control of both outputs in the ratio. For example, ATP/ADP is typically highly sensitive to changes in ADP, and so its regulation is more effective via control of ATP and ADP (or ATP and ATP+ ADP), rather than just ATP. Thus the ability for simultaneous, independent control analysed in Section 2 is a foundation for the regulation quantified in Sections 3 & 4.

In the buffer analysis presented, we have focussed on regulation (i.e. minimising deviations about a set point). However, the controllability analysis discussed in Section 2 is also applicable to control (i.e. changing the set point itself). The ability to simultaneously control different outputs can enable the energy environment to be optimised for different cells or compartments. AK controls the relationship between different ratios via *ATP/ADP* = *K*_*a*_ *ADP*/*AMP*, rather than explicitly controlling the ATP/ADP ratio. For this case, the set point of the ratios can be controlled by changing *K*_*a*_ e.g. by varying *Mg*^*2+*^. This control is in conjunction with the other regulatory mechanisms to jointly control the ratios involving ATP, ADP and AMP.

The quantification methods in this paper determine sensitivities in terms of a few key set points (e.g. ATP concentration) and parameters (e.g. equilibrium constants). While these set points and parameters are themselves dependent upon a larger system, our approach enables biological insights and methods of quantification without requiring us to model and/or experimentally measure all drivers of these set points. For example, we were able to quantify the phosphagen system from experimental measurements of ATP, ADP, AMP, Cr and pCr in muscles during/after exercise and in adipose tissue.

Interestingly, the analysis methods are approximations that are most accurate for small changes, but had good fits for experimental data from exercise that had large disturbances and large changes in metabolites e.g. depletion of creatine phosphate.

Throughout this study, we have shown that multiple energy outputs are simultaneously regulated by different buffering and feedback mechanisms in a synergistic fashion. This finding differs from a general view that cells primarily regulate for one energy output, typically the energy charge or a related AMP-based output [5]. However, our findings are still consistent with this view if buffering is ignored as feedback on its own can only simultaneously control one adenylate energy output.

The quantification of metabolic buffers shows that the ATP/ADP and ATP/AMP ratios are highly sensitive to small changes in ATP when they are at high levels (first reported in [27]). It also shows how strongly the phosphagen system regulates these ratios when the ATP/ADP and ATP/AMP ratios are at high levels. Both creatine phosphate and adenylate kinase have large regulatory effects when the ATP/ADP ratio is highly sensitive to changes, while AMP deaminase has a large regulatory effect when the ATP/AMP ratio is highly sensitive to changes. Interestingly, this observation combined with our recent studies about the interaction of buffering and feedback [17, 16] would indicate pCr should also stabilise glycolytic feedback more strongly at high ATP/ADP and thus enable feedback to be more effective under these conditions. We can also observe in Figure 7 that while the energy charge (the standard energy output) has the ‘steadiest’ output, it is not necessarily the most strongly regulated. In human muscles, the ATP/AMP ratio is significantly more strongly regulated than the energy charge by AMPD, although they are more comparable in mouse adipocytes.

In control theory, the simultaneous, independent control of multiple coupled outputs is referred to as ‘multivariable control’ (see also Box 1), which is a topic that has been extensively studied in non-biological contexts [39] but not in biological ones. In this paper, we provide a foundational case study for multivariable control and cellular biology. As the methods are generic and both buffering and feedback are ubiquitous in biology, we believe that this topic will have significantly wider applicability than the case study provided.

## 6 Conclusion

In this paper, we used control theory and novel buffer analysis for open systems to study buffering in energy metabolism. This enabled us to both uncover underlying principles in metabolic regulation and to quantify the effects of critical regulatory mechanisms. We showed the importance of adenylate kinase along with AMP deaminase and its regeneration for simultaneous, independent control of ATP and the adenylate energy ratios. We also quantified their effect and showed that AK regulates the ATP/ADP ratio while AMP deaminase regulates the ATP/AMP ratio and the energy charge. Similarly, we quantified the effect of creatine phosphate, and showed that it is significantly more effective when the ATP/ADP ratio is highly sensitive to changes. The results were shown to be consistent with experimental data for human muscle and mouse adipocytes.

These results reveal a fundamental role of buffering in biological regulation: enabling the simultaneous, independent control of multiple coupled outputs, which adds to its known role of regulating ‘fast’ disturbances and stabilising feedback. These results also reveal that different buffers carry out different roles - in the phosphagen system, AMP deaminase and adenylate kinase carry out the former role, while creatine phosphate and adenylate kinase (to a much lesser extent) carry out the latter roles.

## Author Contributions

Conceptualization, E.J.H., and J.A.; Methodology, E.J.H., and J.A.; Formal Analysis: E.J.H., and J.A.; Investigation, E.J.H., and J.K; Writing - Original Draft, E.J.H.; Writing - Review & Editing, E.J.H., J.A., and J.K; Funding Acquisition, E.J.H.; Supervision, E.J.H. and J.A.

## Acknowledgements

The authors would like to thank David James, Antonis Papachristodoulou and Lake-ee Quek for their comments and suggestions. EJH was would like to thank the contribution of Judith and David Coffey. JRK is supported by the Australian Diabetes Society Skip Martin Early-Career Fellowship.

The authors declare that there are no competing interests.

## Data Availability

All data generated or analyzed during this study are included in this published article and its supplementary information files.

## Appendix

### Reaction modelled for Controllability Analysis of the Phosphagen System

The reaction used to study controllability with adenylate kinase or creatine kinase are

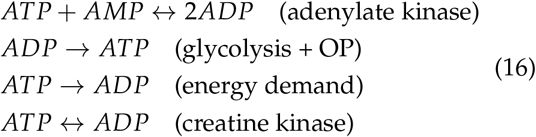

where OP is oxidative phosphorylation. For simplicity, we do not include creatine or creatine phosphate in the CK reaction.

The reaction used to study controllability with AMP deaminase and regeneration are (see Figure 3b)

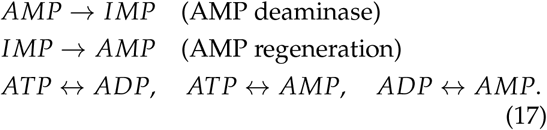

For simplicity, we do not include NH_4_ in the AMPD reaction.

### Figures

Data sources for Figures 5 and 7 are A - [32], B-[191, C-[10], D-[22], E-[13], F-[21]. For repeated cases, the first case shows changes during ATP depletion and the second case shows changes during ATP regeneration.

In Figure **6**, (b) shows sensitivity with *K*_*a*_ = 1 and (c) shows sensitivity with *K*_*a*_ *=* 1 and *B*_*a*_ *=* 1.

Simulations of the model (1) in Figure 1 use constant *p, r = γ y*, where *γ =* 0.1, and *g = − b*_*1*_*y* + *b* _*−1*_ *x* where *b*_1_= 500, *b -*_1_ = 100 for the buffering case (resulting in *B =* 5).

Simulations of the model (5) in Figure 3 use *p = u*_*h*_, *r = γ y*_*1*_ and g = **0**,*r − p*, where *u*_*h*_ is a control input (negative unit step) and *γ* is a kinetic constant.

## Supplementary Information

### SI1 Buffer Analysis Background

This section of the Appendix provides background on the buffer equilibrium ratio and sensitivity function as well as their importance in studying temporal dynamics and differential equation models of regulationn

#### SI1.1 General Buffering Models, Linearization and the Buffer Equilibrium Ratio

Consider the model of a single regulated species that is buffered (see Figure 1)

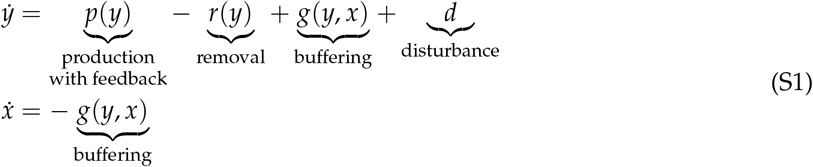

where *y* is an output concentration being regulated, *x* is the concentration of the buffering species, *p* is the production rate of *y, r* is the removal rate of *y, g* is the net flux through the buffering reactions, and *d* is a disturbance to the system. Incorporation of feedback is represented by the *y* dependence of production.

We define the nominal steady state of the system to be its zero-disturbance *(d =* 0) steady state, and denote it as 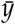, with a corresponding state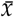; here 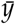 is called the set point of the regulated variable.

The nominal steady state 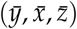 of Equation (S1) occurs when

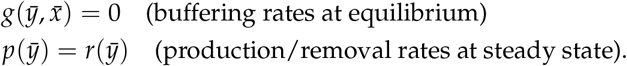

To analyze the model, we reformulate (S1) in terms of deviations 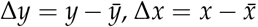, and linearize about the nominal steady state to obtain

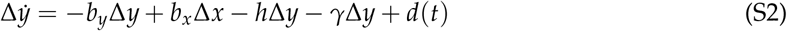

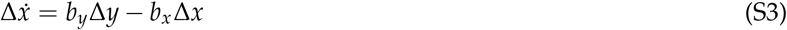

Where

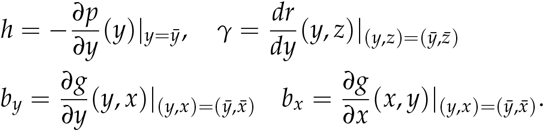

In the main section, we define the buffer equilibrium ratio to be

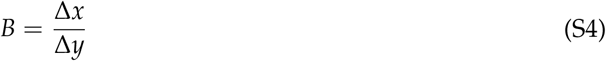

when the buffering reactions are at equilibrium.

In model (S2), if we assume (rapid) equilibrium buffering [17] with quasi-equilibrium of Δy and Δx (by setting 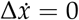 and slow variable Δ*y*_*T*_ *=* Δ*y* + Δx, then we obtain

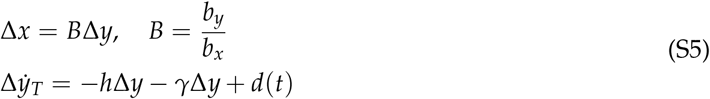

where we can see that (S4) occurs naturally.

Substituting Δ*x* = *B*Δ*y* into Δ*y*_*T*_ *-*Δ*y* + + Δ*x* we have

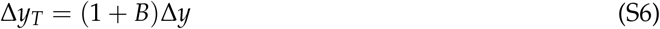

and so

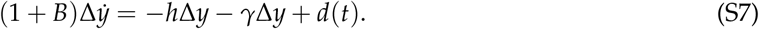

#### SI1.2 Calculating Buffer Equilibrium Ratio B

We can use an explicit or implicit approach to calculate the buffer equilibrium ratio. If we have an explicit function of the buffer concentration *x = x(y)* in terms of output *y* (e.g. assuming buffering at quasi-equilibrium), then we can calculate buffer equilibrium ratio *B* directly using

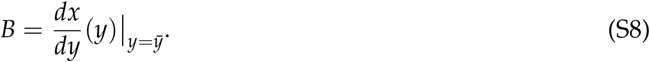

We can also have an implicit relationship between *x* and *y* by setting 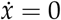 in (SI), where

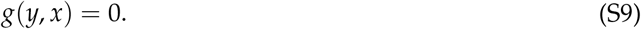

We can use the implicit f function theorem, which results in

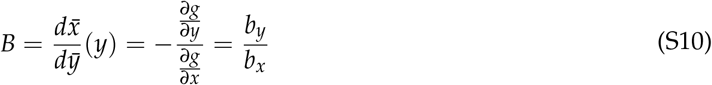

where *b*_*y/*_ *b*_*x*_ and *γ*_*x*_ are defined under (S2).

#### SI1.3 Equivalence of Buffeis to Feedback

PID (Proportioral-Integral-Derivative) feedback controllers are highly important in control theory and are ubiqitous regulatory components of many technological systems [4]. Proportional and integral feedback have been well studied in biological contexts [29, 33, 12, 24, 2, 23, 26, 23, 3, 3]. However, significantly less is understood about the role of derivative feedback. Our recent work showed that rapid buffering is equivalent to derivative feedback and that the buffer equilibrium ratio represents the magnitude of “derivative feedback” [17,16]. This can be seen by noting that with rapid buffering (S7) can be rearranged as

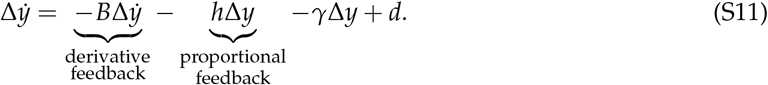

Substituting the linearisation into the net buffering reaction rate, we have the approximation

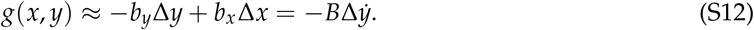

#### SI1.4 Example of a Sensitivity Function

We next calculate the sensitivity function for a simple example with an oscillating disturbances. The sensitivity function is (see Section SI1)

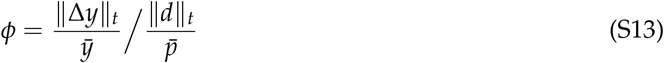

where ‖Δy‖_*t*_ represents the amplitude of oscillations of the output, ‖Δd‖_*t*_ represents the amplitude of oscillations of the disturbance, the output oscillations are normalised by the output set point and the disturbance oscillations are normalised by the steady state production

we have the model

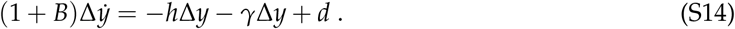

Normalising time in (S7) by the unregulated time-scale 1 /*γ* we have

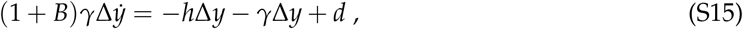

We can use Fourier trarsforms to determine the relative size and phase of oscillating outputs against oscillating disturbances [4]. From (S13),the sensitivity function is

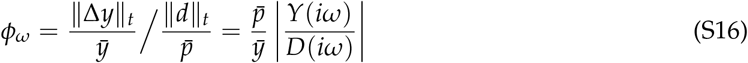

Where *γ* is the Fourier trarsform of *y, D* is the Fourier transform of *d, ω*_*n*_ is the frequency of the oscillations/and | · | is the magnitude of the transform. The transfer furctionfor (SI5) is

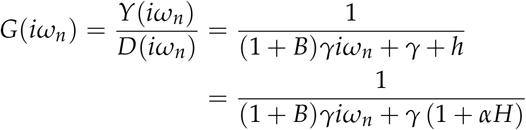

where we set

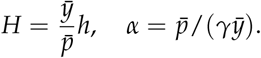

Taking the magnitude of the transfer functions, we have

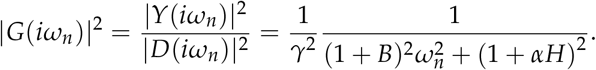

Substituting into (S16) and noting that 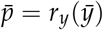, we have

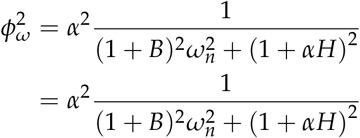

where 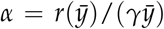 is a measure of the nonlinearity of the removal of *y* [17]. If the removal of *y* is linear then 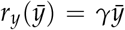 and *α* = 1. We can observe that the sensitivity function is a function of *ω*_*m*_ *H* and *B* and *α*. For the case of no feedback, we have

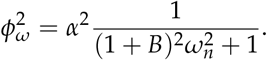

### SI2 Controllability Analysis

In this section, we study the controllability of *ATP, ADP* and *AMP* with buffering and feedback. Controllability is an important system property that determines whether overall regulation and control is possible (see [39,4] for controllability background). Controllability determines the ability for systems inputs (e.g. enzyme activity) to be able to change the states (e.g. metabolite concentrations) of a system.

To model regulatory mechanisms, we assume that the reaction AMP→ADP occurs via adenylate kinase and that regeneration of ATP, ADP and AMP from other sources occurs via the regeneration of AMP from IMP Other regeneration reactions also occur in cells, such as the production of AMP from the degradation of RNA, where if required these reactions can be incorporated in the models as disturbances rather than energy regulatory mechanisms. We do not make assumptions regarding ATP consumption and the models incorporate all forms of these processes. Below, we also analyse the case of polyphosphate kinase, which is an energy buffers that is closely related to the phosphagen system, showing equivalent results.

For clarity of results, we also assume that the reactions all occur in a single compartment. Without this assumption, we would need to model the phosphagen system and corresponding adenine nucleotide concentrations in multiple compartments, which would unnecessarily complicate analysis without providing greater insight.

We first analyse the controllability of a subsystem when studying Adenylate Kinase and Creatine Kinase. We then assume controllability of this subsystem to analyse controllability of the whole system when studying AMPD. This method can be viewed as a form of mathematical induction. Thus, despite using different systems in the following analysis, this approach ensures controllability of the whole system and thus the independent control of multiple outputs. The use of the two systems simplifies analysis by allowing AK and AMPD to be analysed separately.

#### SI2.1 Controllability: Adenylate Kinase

**Figure S1:**
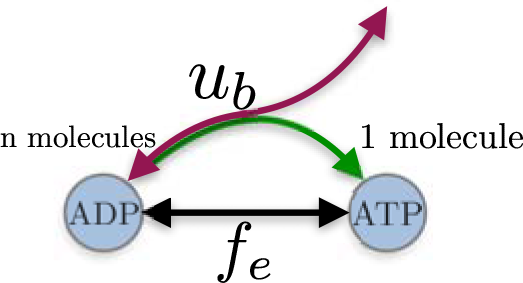
A representation of Model (S17) that shows the buffering flux as a control input *u*_*b*_. The function *f*_*e*_ represents metabolic pathways reactions, including glycolysis, oxidative phosphorylation and ATP demand. The buffer flux represents reactions for creatine kinase *(n =* 1) or adenylate kinase (*n* = 2). The model is shown without the intermediate metabolites *z* that are included in Model (S17). In comparison to the main section, the models treat the buffer as a control input (open loop), rather than a reaction with a rate that is a function of the system concentrations.

Consider the general input-output model of ATP and ADP

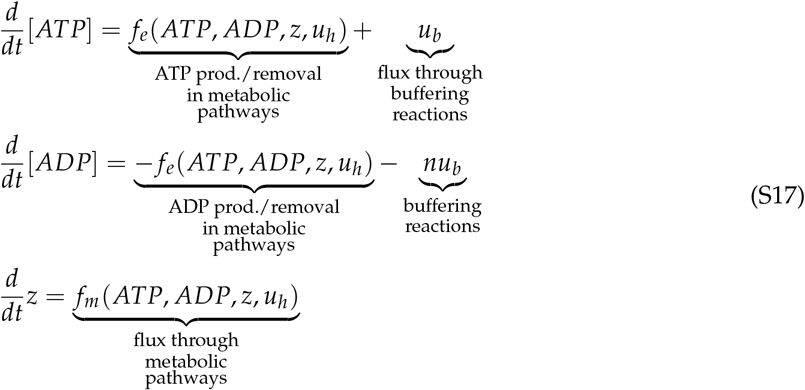

where *z* is an *m* dimensional state representing metabolites excluding ATP and ADP, *f*_*e*_, represents the net flux of all ATP production and removal, *f*_*m*_ is the net flux of all *z* production and removal,*u*_*h*_ is a *p* dimensional input represent the input for feedback control on ATP production, *u*_*b*_ represent the buffering input or buffering flux, and *n* represents the stoichiometry of the buffering reaction *(n = 2* for AK and *n* = 1 for pCr). We ignore *ATP* → *AMP* and *ADP* → *AMP* reactions as they would not be regenerated without adenylate kinase or a similar mechanism. This would result in a stable equilibrium at *ATP* = 0, and so is not useful or informative for controllability analysis.

The steady state 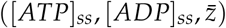 of (S17) occurs when *f*_*e*_*=*0, *f*_*m*_*=o* and *u*_*b*_ =0 for n ≠ 1 and occurs when *f*_*e*_ + *u*_*b*_, = 0 and *f*_*m*_ = 0 for *n* = 1. In the f oil owing, we use the notation 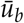 for the steady state of the buffer input.

To simplify controllability analysis, we trarsform the model, such that

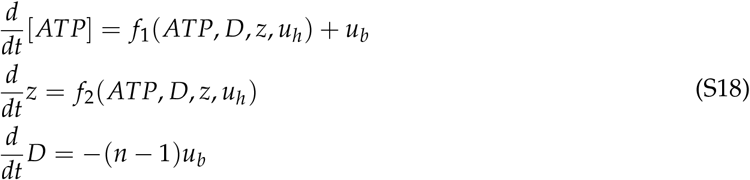

Where *D =* [*ATP*] + [*ADP*],

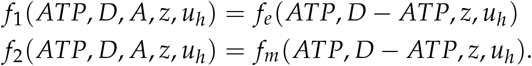

We can observe in Equation (S21) that D is only controllable with.*u*_*b*_ and with *n* ≠1. Without *u*_*b*_ or with *n* = 1 then D is constant. Below shows the same results with a more formal linear controllability analysis.

We use a linearisation of the model for analysis, which is consistent with the linearized approach in all other sections of the paper. Linearising (S21) and placing in a standard control form, we have

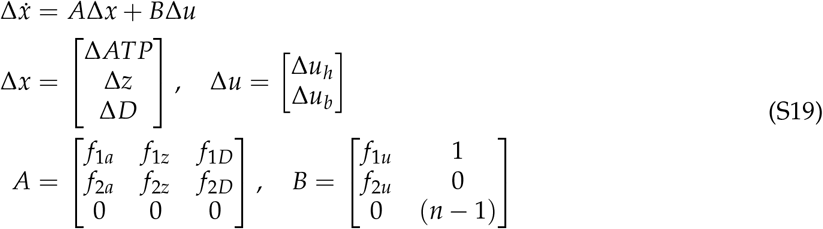

where 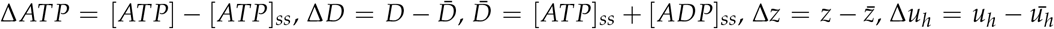 and 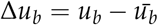 are the deviations of the respective variables from their steady states, and

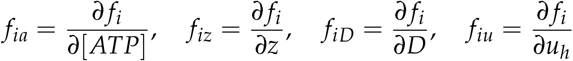

are the linearisations of the functions about the steady states, where f_i_ represents *f*_*1*_ or *f*_*2*_.

The controllability matrix is ζ = [B/AB, A^**2**^B,…, A^m+**2**^B] and the system is controllable if the matrix has a rank of *m* + 2 *[39*,4].

We first note that if there is no buffer control then

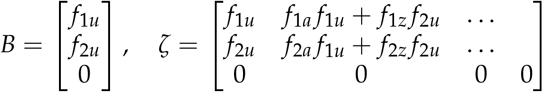

and so the system is not controllable as the matrix rank is less than *m* + **2**.

If there is buffer control then

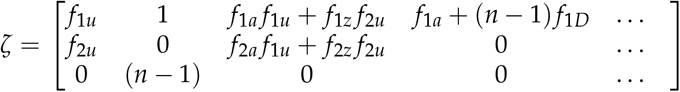

If we assume that the subsystem that excludes D is controllable then the top *m* + 1 rows of ζ have a rank of *m* +1. In this case, the matrix has rank *m* + 2 if *n* ≠ 1, and so the system is controllable for AK (*n* = 2) but not pCr (*n* = 1).

We can obtain an equivalent result for output controllability, where we are only concerned with the controllability the outputs ATP and ADP, but not the intermediate metabolites. This result can be used to remove the assumption on controllability of the intermediate metabolites. The standard output form for this case is

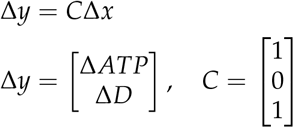

which-along with (SI 9), described the relationship between inputs, states and outputs.

The output controllability matrix is ζ = [*CB,CAB,CA*^*2*^*B*,.. ., CA^m+**2**^B] and the system is output controllable if the matrix has a row rank of 2 [39,4]. If we assume that Δ *ATP* is output controllable, rather than assuming the subsystem without *D* is controllable, then the output controllability follows from the above derivation and the modified controllability matrix.

#### SI2.2 Controllability: Polyphosphate Kinase

Above, we assumed that AMP → ADP occurs via adenylate kinase, which is the dominant path [7]. However, the reaction can also occur through buffering reactions that are closely related to the phosphagen system, such as specific forms of polyphosphate kinase [1], where polyphosphate carries out an analogous role to creatine phosphate in microorganisms. For this case, the results still hold if we group these enzymes with adenylate kinase (see SI2 for analysis), just as we grouped creatine kinase with feedback on metabolic pathways (See Figure 3).

Consider the general input-output model of ATP and ADP

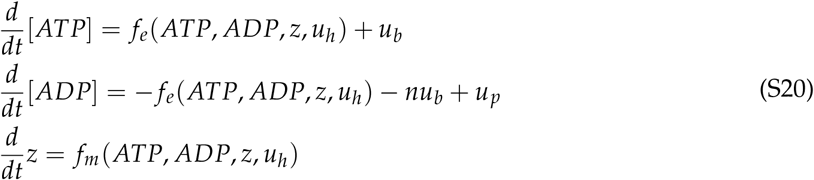

where *u*_*up*_ represents enzymes that can carry out AMP→ ADP To simplify controllability analysis, we once again transform the model, such that

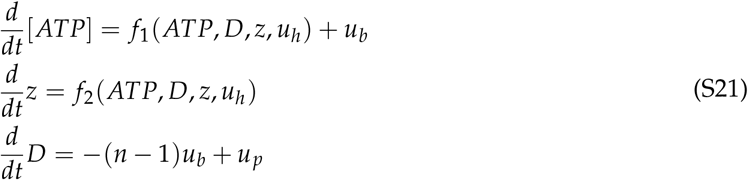

where D = [ATP] + [ADP],

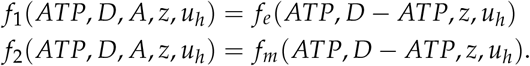

We can observe in Equation (S21) that D is only controllable with either *u*_*p*_ or *u*_*b*_. Without either mechanism then D is constant. Formal linear controllability (or output controllability) analysis can be completed as above with the same results.

#### SI2.3 Controllability: AMP Deaminase

**Figure S2:**
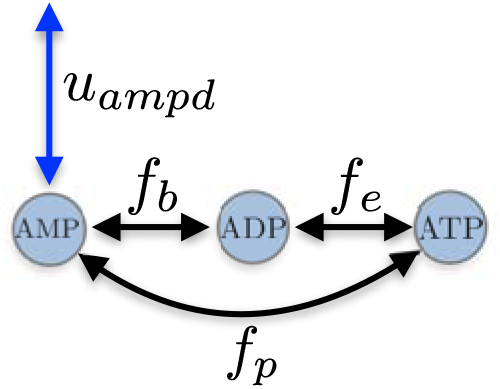
Representation of Model (S22). Shown without intermediate metabolites z.

We next look at the case for AMP deaminase. Consider the general input-output model of ATP, ADP and AMP

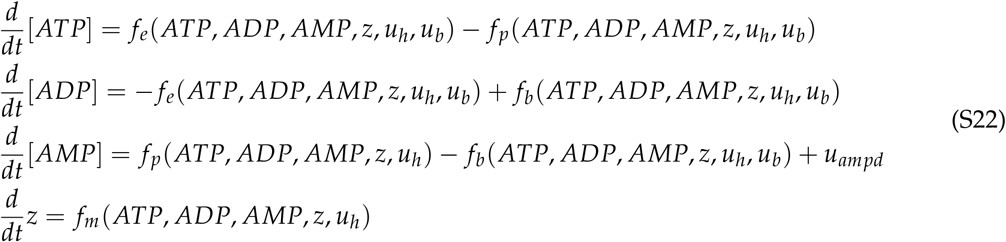

where *z* is an *m* dimensional state representing metabolites excluding ATP, ADP and AMP,*u*_*b*_ represent the buffering input (e.g. Adenylate Kinase) and *u*_*ampd*_ represents the buffering input from AMP deaminase. represents the net flux of all ATP ↔ *ADP, f*_*p*_ represents the net flux of all *ATP* ↔*AMP* and *f*_*b*_, represents the net flux of all *ADP* ↔ *AMP*.

The steady state 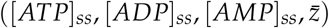 of (S22) occurs when *f*_*m*_ *-* 0, *= u*_*ampd*_ -*0, = f*_*e*_ *- f*_*p*_ In the foil owing, we use tine notation 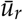 for the steady state of the AMPD input.

To simplify controllability analysis, we trarsform the model, such that

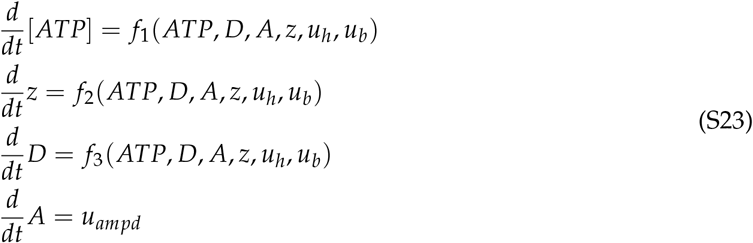

where *D* = [*ATP*] + [*ADP*], A = *[ATP]* + [ADP] + [AMP], and

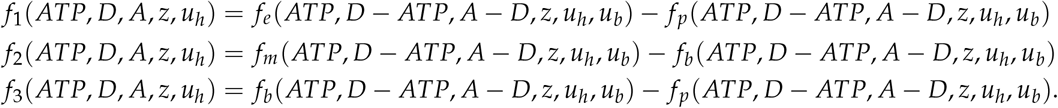

We can observe in (S23) that *A* is only controllable with *u*_*ampd*_. Without *u*_*ampd*_ then *A* is constant. Formal linear controllability analysis can be completed as above with the same results, both for standard or output controllability.

Thus the energy metabolism system is only controllable with both AK and AMP deaminase. Further, without AK and AMPD then feedback on metabolic pathways is fundamentally limited to only controlling one combination of ATP, ADP and AMP

#### SI2.4 Controllability: Equivalence with Energy Ratio Output Coordinates

We next show that controllability of the system is equivalent with different outputs. Controllability is analysed with the outputs (y, *X*_*d*_, *A)* where

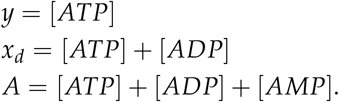

This can be rewritten as

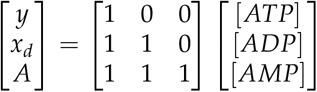

where the matrix is invertible and so there is a unique mapping between the *(y, x*_*d*_, *A)* and the individual concentrations.

We next show that controllability of the system with outputs *ATP, ADP* and *AMP* is equivalent to controllability of the system with outputs *ATP, ATP/ADP* and *ATP/AMP*, where the relationship is nonlinear. The second set of outputs *(y,y*_*rd*_, *y*_*rm*_) are

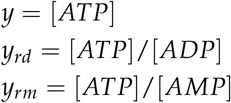

For positive concentrations there is a unique mapping between the two output coordinates resulting from the relations

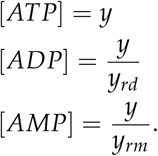

Thus controllability in one set of output coordinates is equivalent to controllability in the other set of output coordinates.

### SI3 pH Buffering in Open Systems

The reader may find it instructive to compare and contrast the open-system metrics introduced in Section SI1 to familiar metrics commonly used to analyze closed-system buffering. For ease of readability, we describe the results in SI3 and include the working in SI4.

Consider the well-studied application of pH buffering, and more specifically, the simple example of a weak acid buffer. Such a system can be represented by the reaction

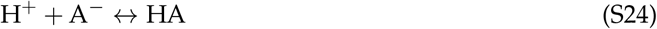

where HA is a weak acid that buffers H^+^ (and thus pH, given that pH = − log_**10**_[H^+^]),and the thermodynamic equilibrium constant *K =* [H^+^][A^-^]/[HA]. (Here, we ignore the water ionization reaction) For reaction (S24), the buffer equilibrium ratio *B* is (see SI4.1)

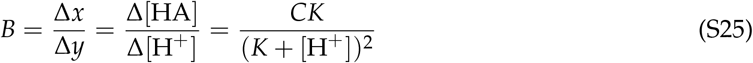

where the buffer concentration is *x =* [HA], the output concentration is y = [H^+^], C = [HA] + [A^−^], and 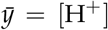 is at the output set point. Figure S3 demonstrates that *B* is a function of [H^+^] and £, increasing with decreasing [H^+^]. If [H ^+^] is fixed, *B* takes on a maximum when *K* = [H^+^].

**Figure S3:**
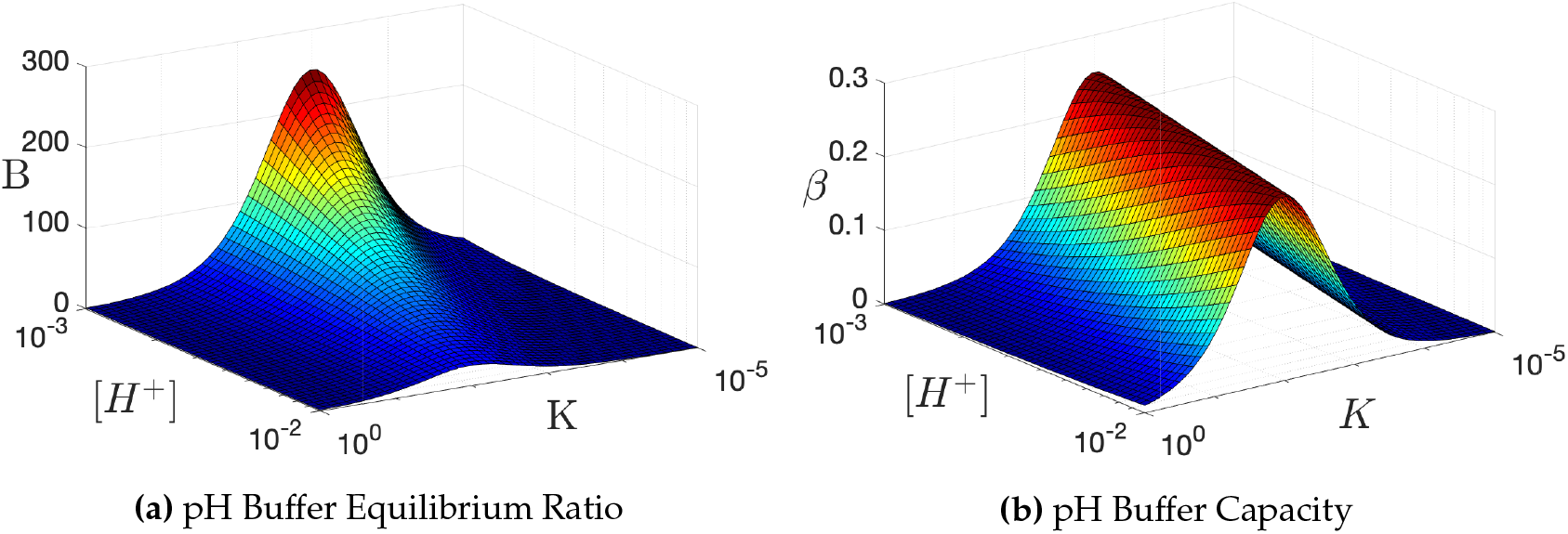
pH buffer capacity (bottom) and buffer equilibrium ratio (top) with *C* = [*A*^−^] + [*HA*] = 1. *β* has concentration units while *B* is dimensionless.

In pH theory, the buffer capacity metric *β* = Δ/ΔpH, is a measure of the resistance of a buffer solution to pH change on addition of a strong base *(n* represents the moles of added OH ^−^ ions) [40]. The sensitivity function, *ϕ*, is its (inverted) open-system generalization [40]:

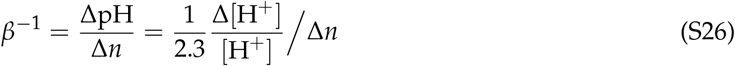

where [H^+^] = *y* is the output signal being regulated. The form of (S26) is similar to (4) with a few exceptions: (a) the 2.3 factor, which results from the log(10) scaling of pH; (b) the disturbance, Δ[H^+^] can be interpreted as a one-time addition of mass (OH^−^) to an otherwise closed system, whereas the sensitivity function, ϕ, considers a system where the disturbance, *d*, is directly on the rate of change of the output signal; and (c) the disturbance, Δ[H^+^], is not normalized by a production term, as there is no on-going rate of H^+^ production in the closed system.

Note that in the closed pH buffering system, both *B* and *β* convey the isolated effectiveness of the buffer. In fact,

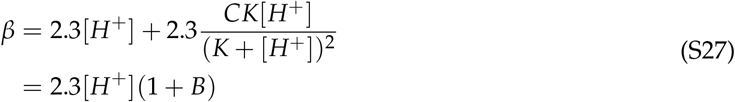

(Furthermore, the constant 2.3[*H*^+^] term is often ignored, as it typically achieves significance at pH levels well below levels of interest [40].) In the open systems, whilst *B* still represents an isolate measure of the buffer, the sensitivity function, *ϕ*, depends upon the type of disturbance and other forms of regulation present [16,17] (see SI1.4 for a simple example).

The plot of the buffer capacity *β* can be seen in Figure S3. If [*H* ^+^] is fixed, then buffer capacity is maximum when *K* = [*H* +], similar to *B*. However, if [*H* ^+^] is varied then is maximum when [*H* ^+^] = *K*, rather than a maximum of *B* at small [*H* ^+^]. This change results from the buffer capacity including an extra [*H* ^+^] for normalisation, which in open systems is included in the sensitivity function (4) rather than *B*.

### SI4 Buffer Equilibrium Ratios

This section of the Appendix provides supporting evidence of buffer equilibrium ratio calculations presented in the main section of the paper. We determine the buffer equilibrium ratios for different reactions.

#### SI4.1 pH Buffering

The first reaction that we study is the pH buffering reaction

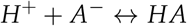

which can be written as

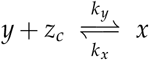

where *y* = [*H*^+^], *x* = [*HA*] and cosubtrate *z*_*c*_ *= [A*−]. The forward and reverse reaction rates are

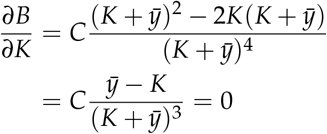

and at equilibrium we have

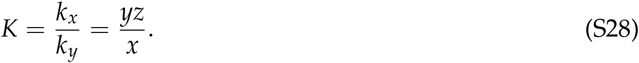

Setting C = *z*_*c*_ + *x* and rearranging, we have

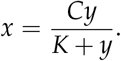

The buffer equilibrium ratio can be calculated as

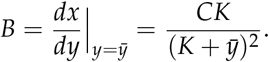

Substituting 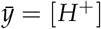 then buffer equilibrium ratio is

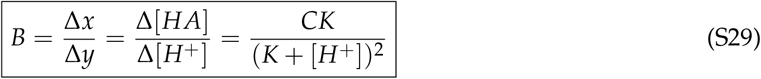

which is shown in (S24) in SI3.

##### SI4.1.1 Optimal Parameters

For fixed kinetic constants, B is maximum when 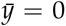, for which *B* = *C/K. B* approaches zero when 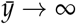. The buffer equilibrium ratio *B* can also be increased linearly by increasing C.

For a fixed non-zero 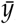 the maximum buffer equilibrium ratio occurs when

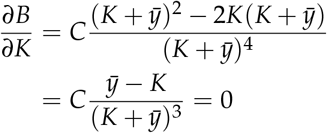

which occurs when

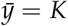

and results in

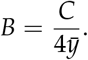

For a fixed nonzero *K*, the maximum buffer equilibrium ratio occurs when 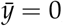, and results in

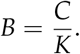

#### SI4.2 Creatine Phosphate

The next reaction that we study is the creatine kinase reaction

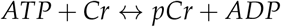

where Cr is creatine and *pCr* is the creatine phosphate that buffers *ATP*. We can rewrite this in the form

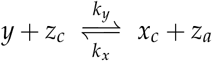

where *z*_*c*_ = [*Cr*], *x* = [*pCr*], *y* = [*ATP*] and *z*_*a*_ = [*ADP*].

##### SI4.2.1 Buffer Equilibrium Ratio: Constant Parameters

The forward and reverse reaction rates are

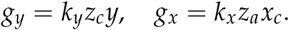

and at equilibrium we have

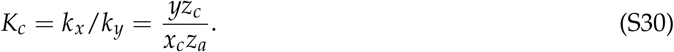

Setting *C = z*_*c*_*+x* and *D =y+z*_*a*_ and rearranging, we have

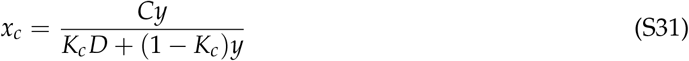

The Buffer Equilibrium Ratio for fixed parameters is

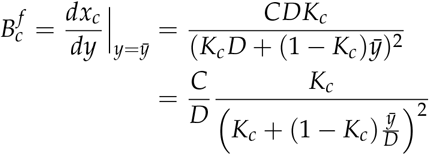

which is a function of the three independent inputs 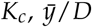 and C/D. Substituting 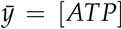, we have

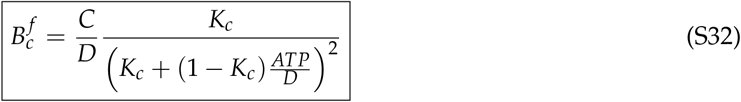

which is shown in (7) in the main section.

##### S 14.2.2 Buffer Equilibrium Ratio: Varying parameters *K*_*c*_ and *A = ATP* + *ADP*

If *K* and *D* in (S31) vary with ATP then we need to incorporate varying parameters into the buffer equilibrium ratio *B*_*c*_. From (S31), we have the buffer concentration

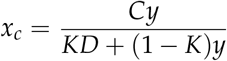

and so

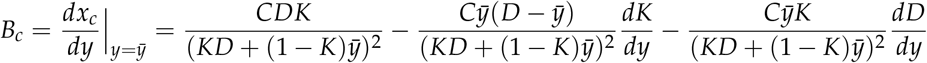

where the derivatives of i<and D are evaluated at 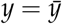. Using the notation 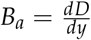 (see Section SI4.3 for further explanation) then

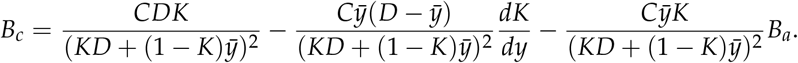

Using (S32), we set

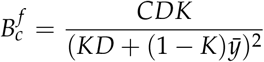

and so

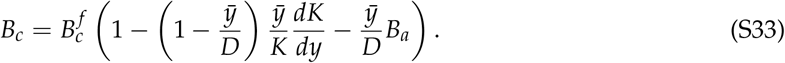

Substituting 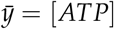, we have

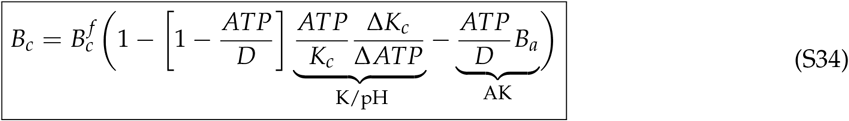

which is shown in (**11**) in the main section.

Equilibrium constant *K*_*c*_ typically increases with decreasing ATP (via decreasing pH), where *ΔK*_*C*_*/ΔATP* < 0 (see SI4.2.3). Thus the effect of changing *K*_*c*_ is to increase *B*_*c*_ in (11). In contrast, *D* decreases with decreasing ATP *(B*_*a*_ > 0) and so the effect in (11) of changing *D* is to reduce *B*_*c*_. *As* such, adenylate kinase reduces the effectiveness of buffering by creatine phosphate in (**11**).

##### SI4.2.3 Comparison to Experimental Data

We next compare the theoretical and experimental values f or *B*_*c*_ in human muscles and adypocyte. To determine the experimental *B*_*c*_, observations of *pCr* and *ATP* over time (e.g. before and after a period of exercise’] can be used in the formula

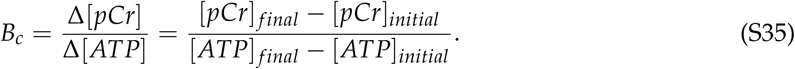

This formula is a simple method of calculation, but can result in larger measurement errors for large *B* if there are small changes to [ATP].

To calculate the theoretical B over a range of *y*, we use

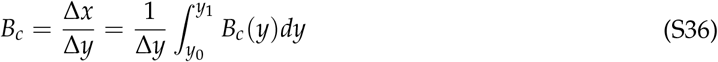

where *Δy – y*_*1*_ *− y*_*0*_. We know experimental values of *k*_*c*_., C/D and *y/D* for particular measurement points rather than all values, and so for fixed *B* we approximate the integral using the trapezoidal rule

**Table S1:** Buffer equilibrium ratio *B*_*c*_ of creatine phosphate for human muscle during and after exercise. *not shown in Figure *5* due to large experimental error.

| $B_{exp}$ | $B_{theor}$ | $B_{theor}$<br>inc. $\frac{dK}{dy}, \frac{dD}{dy}$ | $K_{exp}$ | $\frac{ATP}{ATP+ADP}$<br>(exp) | Ref |
| --- | --- | --- | --- | --- | --- |
| 11.5±8.6 | 5.8 | 12.6 | 4.9-190 | 0.88-0.83 | [32](ATP ↓) |
| 15.4±5.8 | 6.7 | 15.2 | 2.5-15.4 | 0.87-0.85 | [19](ATP ↓) |
| 20.9±9.9 | 7.1 | 19.0 | 15.4-2.6 | 0.85-0.87 | [19] (ATP ↑) |
| 7.5±2.2 | 5.5 | 8.4 | 17.9-138.8 | 0.87-0.82 | [10](ATP ↓) |
| 10.7±3.6 | 5.5 | 10.8 | 13.9-86.9 | 0.86-0.77 | [10](ATP ↓) |
| 7.3 ±1.5 | 7.7 | 8.2 | 5.8-76.5 | 0.90-0.79 | [22](ATP ↓) |
| 16.0±9.9 | 11.42 | 16.3 | 76.5-3.6 | 0.79-0.89 | [22](ATP ↑) |
| 8.3±1.5 | 8.3 | 7.6 | 4.6-42 | 0.92-0.87 | [13] (ATP ↓) |
| 36±49.5 * | 9.6 | 35.6 | 42-5.4 | 0.87-0.89 | [13] (ATP ↑) |

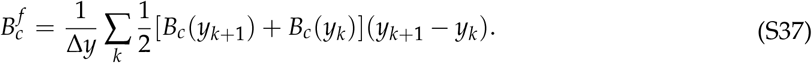

The theoretical 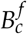 is estimated from the experimental *K, C* and *D* at each data point.

We can use a similar approach for the case with varying parameters, where we average the fixed components and the *K* and *D* rate of change components. When applying the average for this case, we use the geometric average as the parameters can vary over orders of magnitude (see Table S1). We continue to use the arithmetic average for the fixed component for consistency with above. If we set

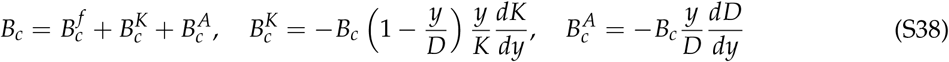

then we use the approximation

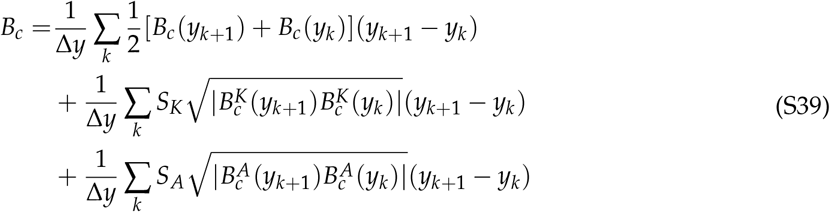

where *S*_*K*_ = sign(*BK*(*y*_*k*+1_)*B*_*K*_(*y*_*k*_)) and *S*_*A*_ = sign(*B*_*A*_(*y*_*k*+1_)*B*_*A*_(*y*_*k*_)).

#### SI4.3 Adenylate Kinase

We next study the reaction adenylate kinase

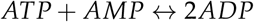

that can be written in the general form

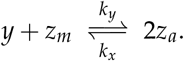

We set

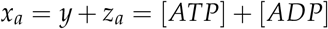

where the demand reaction *y* → *Z* _*d*_does not affect the buffer *x*_*a*_ for this buffer definition.

We note that

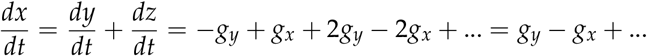

and so the effect of the buffering reactions is equivalent to the buffering in the general model (S1).

We next derive the buffer equilibrium ratio using this definition of the buffer variable.

The forward and reverse reaction rates of

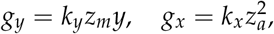

and the equilibrium is

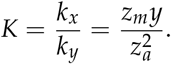

For A= *y*+ x_*a*_,we have

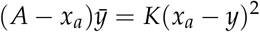

resulting in

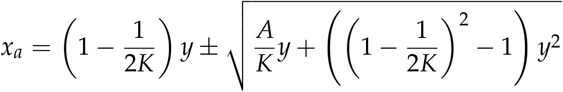

As *x*_*a*_ *− y* ≥ 0, this simplifies to

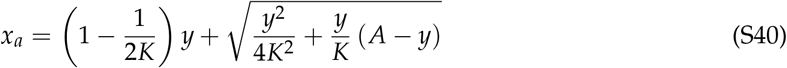

#### SI43.1 Fixed parameters *K* and *A*

Differentating (S40), we have

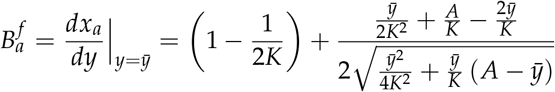

Substituting from (S40), we have

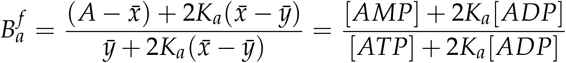

where *[ATP], [A DP], [AMP]* are evaluated at the set point. Thus we have

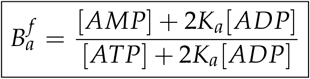

which is shown in (13) in the main section.

#### SI4.3.2 Varying parameters *K* and *A*

We next wish to determine the effect of varying *K* and *A* on buffer equilibrium ratio *B*. From (S40), we have

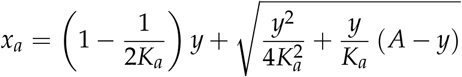

For varying A, we have

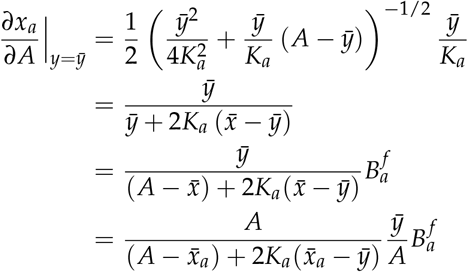

Where

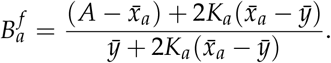

This results in

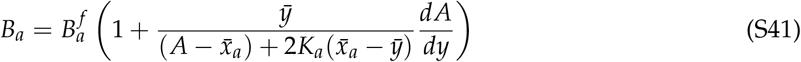

(S42)

where *d A/dy* is evaluated at 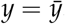. This can be rewritten

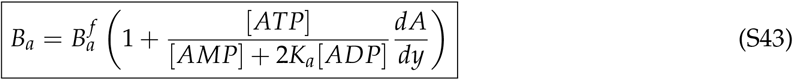

which is shown in (14) in the main section.

The adenine nucleotide total. *A*, typically decreases with decreasing ATP (*R*_*m*_ > 0) and so the effect of AMPD changing A is to synergistically increase *B*_*a*_ in (11).

If we also have *K* that varies with *y* then we can also include extra terms in *B*_*a*_. We have

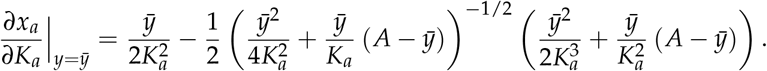

Substituting, we have

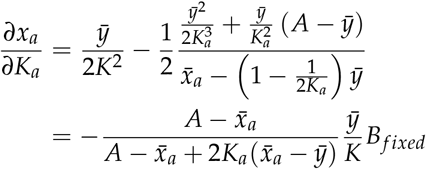

which can be rewritten

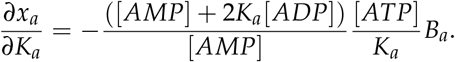

Combining with (S41), we have

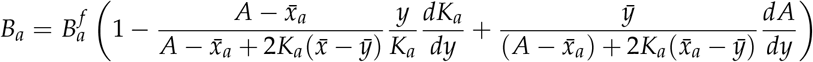

where *d A/dy* and *dK/dy* are evaluated at 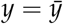. This can be rewritten

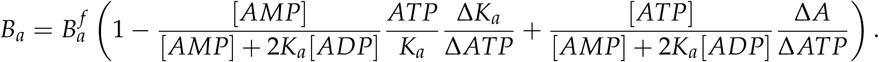

##### SI4.3.3 Comparison to Experimental Data

We next compare the theoretical and experimental values for *B*_*a*_ in human muscles and adypocyte during exercise. To determine the experimental *B*_*a*_, observations of ATP and *ADP* over time (e.g. before and after a period of exercise) can be used in the formula

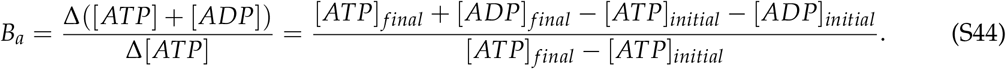

To calculate the theoretical *B* over a range of y, we use

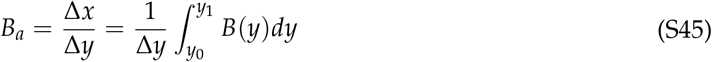

where *Δy = y*_*1*_ *− y*_*0*_. We know experimental values of *K*_*a*_, *y/D* for particular measurement points rather than all values, and so for fixed parameter 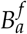 we approximate the integral using the trapezoidal rule

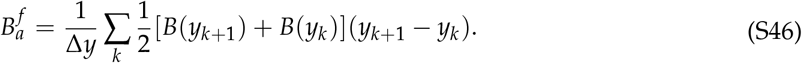

The theoretical *B* is estimated from the experimental *K*_*a*_, *y/D* at each data point.

As above, we use the geometric average as parameters can vary over orders of magnitude. If we set

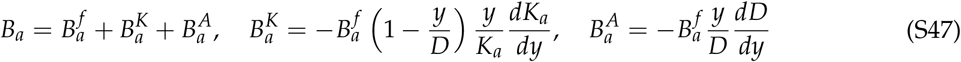

then

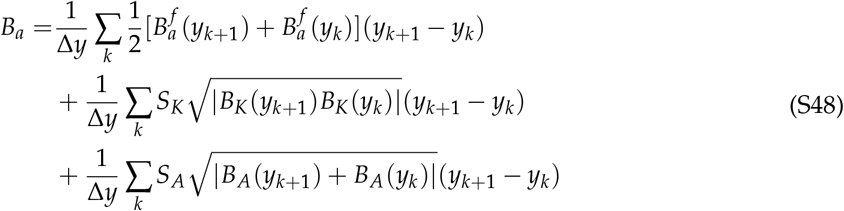

where *S*_*K*_ *=* sign(*B*_*K*_(*y*_*k*+1_) *B*_*K*_(*y*_*k*_)) and *S*_*A*_ *=* sign(*B*_*A*_(y_k+**1**_)*B*_A_(y_k_)).

**Table S2:** Buffer equilibrium ratio *B*_*a*_ of adenylate kinase for mouse and for human muscle during and after exercise. ^*^also incorporates 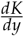.

| $B_{exp}$ | $B_{theor}$ | $B_{theor}$<br>inc. $\frac{dA}{dy}$ | Ref | Tissue |
| --- | --- | --- | --- | --- |
| 0.94±0.09 | 0.11 | 0.96 | [32](ATP ↓) | muscle |
| 0.94±0.05 | 0.05 | 0.94 | [19](ATP ↓) | muscle |
| 0.85±0.09 | 0.05 | 0.84 | [19](ATP ↑) | muscle |
| 0.85±0.27 | 0.13 | 0.87 | [10](ATP ↓) | muscle |
| 0.88±0.41 | 0.09 | 0.88 | [10](ATP ↓) | muscle |
| 0.89±0.07 | 0.08 | 0.90 | [22](ATP ↓) | muscle |
| 0.74±0.17 | 0.05 | 0.76 | [22](ATP ↑) | muscle |
| 0.88±0.10 | 0.44 | 0.89* | [21](ATP ↓) | adipose - basal |
| 0.78±0.42 | 0.44 | 0.79* | [21](ATP ↑) | adipose - basal |

### SI5 Sensitivity Functions and Energy Ratio Regulation

In this section, we determine the sensitivity of the ATP/A*DP ratio, the ATP, AMP ratio and the energy charge to changes in ATP. We also compare these sensitivities to experimental data.

#### SI5.1 Sensitivity function: ATP/ADP Ratio

We next determine the sensitivity of the ATP/ADP Ratio to changes in A*TR We have the output

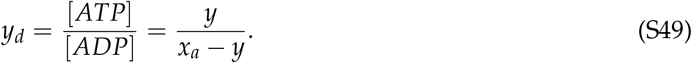

Differentiating, we have

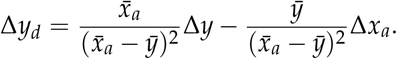

where the derivatives are evaluated at the steady state 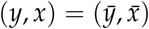 Using *Δx*_*a*_ *= B*_*a*_*Δy*_*f*_ we have

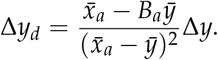

Normalising, we have

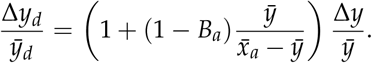

Rewriting, we have

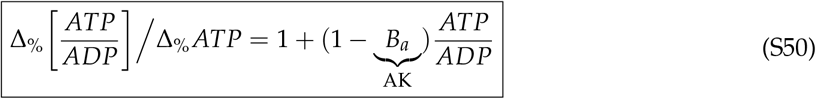

which is shown in (15) in the main section.

#### SI5.2 Sensitivity function: ATP/AMP Ratio

We next determine the sensitivity of the ATP/AMP Ratio to changes in ATP. We have the output

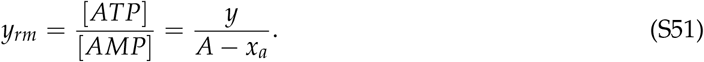

Differentiating, we have

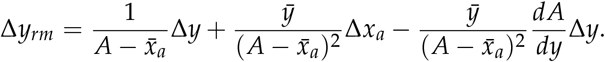

where the derivatives are evaluated at the steady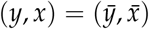. Using Δ *x*_*a*_ = BaΔ *y*we have

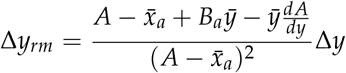

Normalising, we have

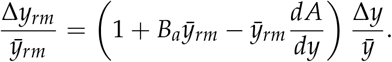

Setting 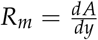 and rewriting, we have

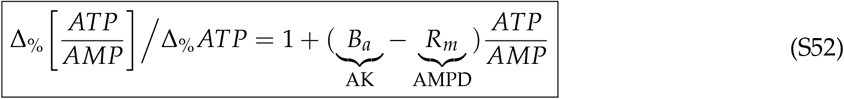

which is shown in (15) in the main section.

#### SI5.3 Sensitivity function: Energy Charge

We next determine the sensitivity of the the Energy Charge Ratio to changes in *[ATP]*. The energy charge can be written

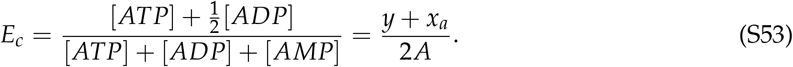

Differentiating, we have

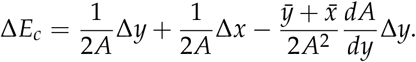

where the derivatives are evaluated at the steady state 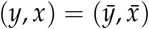. Using *Δx = B Δy*, we have

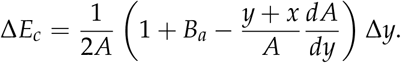

Normalising, we have

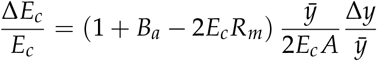

Where 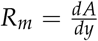, Rewriting, we have

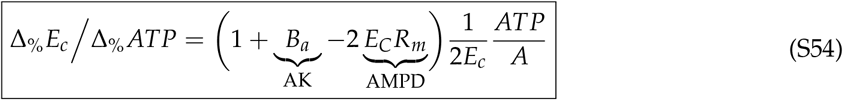

which is shown in (15) in the main section.

#### SI5.4 Comparison to Experimental Data

We can compare the theory to experimental results, which can be seen in the main section For an initial and final value of *ATP, ADP, AMP*, we have

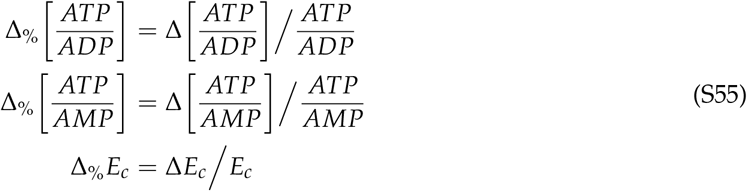

where

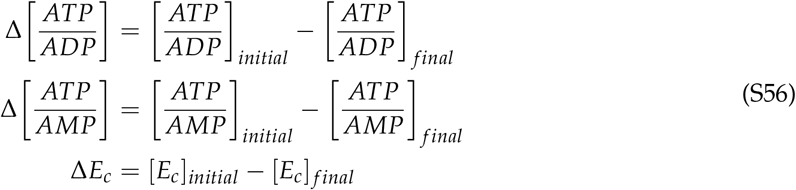

and for the normalisation we use the geometric averages

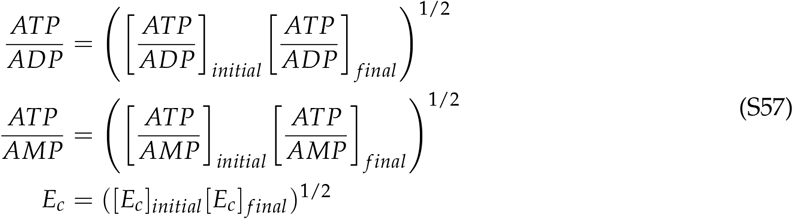

To determine the theoretical results, we substitute the geometric averages of the experimental values for *B*_*a*_, *R*_*m*_, ATP/ A *DP*, ATP/AMP, and *E*_*c*_ into Equations (S50), (S52) and (S54).

#### SI5.5 Comparison of Sensitivity Functions

We can note the difference between the single output sensitivity function (4) described in Section SI1 and the sensitivity functions (15) described in Section 4 that relate different outputs. Equation (4) represents the sensitivity between a disturbance d (e.g. varying flux) and an output y (e.g. ATP), while (15) represents the sensitivity between the ATP concentration and various outputs ratios (ATP/ADP, ATP/AMP and the energy charge). Comparing the sensitivity of the different outputs directly is important in its own right e.g. it is analogous to relating the charge (ATP) and voltage (ratios) of a battery. However (15) and (4) can be used together, where (15) enables the conversion of (4) from one choice of output (e.g. ATP) to another (e.g. ATP/ADP). This conversion allows the effect of a disturbance on different outputs to be analysed. An example conversion can be observed for the sensitivity function in (4’) with outputs:

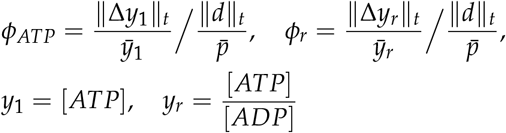

where *d* is the disturbance of interest. We have

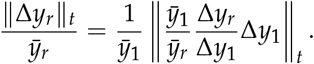

For constant 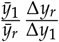, determined at each set point (e.g. ATP) and constant (e.g. *B*_*a*_), then

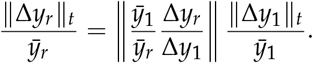

The conversion of (4) between outputs follows as

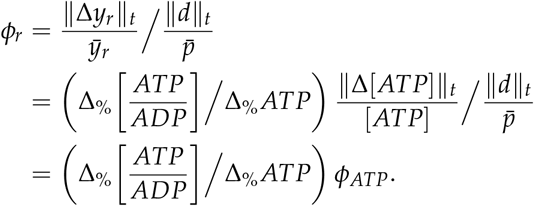

